# MaizeCODE reveals bi-directionally expressed enhancers that harbor molecular signatures of maize domestication

**DOI:** 10.1101/2024.02.22.581585

**Authors:** Jonathan Cahn, Michael Regulski, Jason Lynn, Evan Ernst, Cristiane de Santis Alves, Srividya Ramakrishnan, Kapeel Chougule, Sharon Wei, Zhenyuan Lu, Xiaosa Xu, Jorg Drenkow, Melissa Kramer, Arun Seetharam, Matthew B. Hufford, W. Richard McCombie, Doreen Ware, David Jackson, Michael C. Schatz, Thomas R. Gingeras, Robert A. Martienssen

## Abstract

Modern maize was domesticated from *Teosinte parviglumis*, with subsequent introgressions from *Teosinte mexicana*, yielding increased kernel row number, loss of the hard fruit case and dissociation from the cob upon maturity, as well as fewer tillers. Molecular approaches have identified several transcription factors involved in the development of these traits, yet revealed that a complex regulatory network is at play. MaizeCODE deploys ENCODE strategies to catalog regulatory regions in the maize genome, generating histone modification and transcription factor ChIP-seq in parallel with transcriptomics datasets in 5 tissues of 3 inbred lines which span the phenotypic diversity of maize, as well as the teosinte inbred TIL11. Integrated analysis of these datasets resulted in the identification of a comprehensive set of regulatory regions in each inbred, and notably of distal enhancers which were differentiated from gene bodies by their lack of H3K4me1. Many of these distal enhancers expressed non-coding enhancer RNAs bi-directionally, reminiscent of “super enhancers” in animal genomes. We show that pollen grains are the most differentiated tissue at the transcriptomic level, and share features with endosperm that may be related to McClintock’s chromosome breakage-fusion-bridge cycle. Conversely, ears have the least conservation between maize and teosinte, both in gene expression and within regulatory regions, reflecting conspicuous morphological differences selected during domestication. The identification of molecular signatures of domestication in transcriptional regulatory regions provides a framework for directed breeding strategies in maize.

## Introduction

Modern maize (*Zea mays* ssp. *mays*) is the result of domestication from its ancestor *teosinte parviglumis* (*Zea mays* ssp. *parviglumis*), with subsequent introgressions from *teosinte Mexicana* (*Zea mays* ssp. *mexicana*)^1–6^. Domestication traits include increasing the number of kernels per ear, limiting tillering, removing the hard fruitcase and preventing the kernels from disarticulating upon maturation^3^. The genetic study of the domestication process has led to the identification of many key regulators, mostly transcription factors, responsible for some of these traits. The most important of these regulatory genes is t*eosinte branched 1 (tb1)*, which encodes a TCP transcription factor (TF) with a basic helix-loop-helix DNA binding domain^7,8^. TB1 is a master-regulator, regulating other TFs as well as itself, in a tissue-specific manner^8,9^. Among its targets, *grassy tillers 1 (gt1)* promotes apical dominance along with TB1 in modern maize^10^. Several other genes have been implicated in the domestication or improvement processes, such as *tunicate 1 (tu1)*^11,12^, *ramosa1 (ra1)*^13^ and *teosinte glume architecture (tga1)*^14^ but it is now clear that a complex epistatic network of genes has evolved through domestication, which relies more on quantitative regulation than presence/absence^2–4^.

Identification of regulatory regions has been pioneered in animals by the ENCODE (Encyclopedia of DNA elements) project^15,16^. ENCODE relies on integrating datasets which evaluate chromatin structure, such as measuring DNA accessibility and histone post-translational modifications, with transcriptomic datasets to identify the patterns that could regulate and/or register gene expression. In animals, regulatory regions, also called enhancers, are usually marked by mono-methylation of lysine 4 on the histone H3 tail (H3K4me1), with active enhancers showing acetylation of other lysines on the H3 tail (H3ac) and inactive enhancers having H3K27me3^17^. Clusters of regulatory regions have additional signatures, notably the presence of capped RNA molecules called enhancer RNAs, which are transcribed from both strands of DNA in these distal regulatory regions.

The study of cis-regulatory regions in plants have revealed similar molecular signatures^18^, although H3K4me1 is found in gene bodies rather than distal enhancers^9,19,20^, and the presence of enhancer RNAs is disputed^21^. The catalog of distal elements in maize has been greatly improved by recent efforts to resolve cell-type specific accessible regions with single-cell ATAC-seq experiments^9,19,22^. ATAC-seq identifies nucleosome free and other open chromatin regions, which include many but not all promoters and enhancers, as well as many other regions of the genome accessible to bacterial transposase. While ATAC-seq is uniquely powerful in the single cell context, open chromatin alone does not generate a comprehensive dataset of regulatory regions^23^, especially when limited to selected tissues and inbred lines.

By careful selection of tissues, inbred lines, and epigenomic signatures we present MaizeCODE, a comprehensive catalog of maize cis-regulatory regions, accompanied by a computational pipeline for their analysis. We also present datasets from the teosinte inbred line TIL11, as well as a chromosome level genome sequence assembly, allowing us to investigate the impact of domestication on gene regulation. Through this integrated analysis, we identified tissue-specific enhancers with bi-directionally expressed enhancer RNAs, in each tissue of all inbreds. These “super-enhancers” are more accessible to regulatory TFs and inherently drive higher transcription levels. Interestingly, “super-enhancers” also have stronger RdDM signals at their boundaries, including both polIV and polV transcripts. This could reflect the co-evolution or co-regulation of active regulatory regions and the silencing of neighboring TEs. We illustrate the utility of these datasets and their analysis by demonstrating the tissue-specific impact of domestication on the conservation of enhancers and of the genes they regulate. For example, we demonstrate that regulation of ear development was a major target of maize domestication. We also uncover variation in telomere maintenance in pollen and endosperm that could underlie McClintock’s chromosome breakage-fusion-bridge cycle.

## Results

### Reference genome assemblies and data types selected for MaizeCODE

We selected one stiff-stalk (B73), one non-stiff-stalk (W22) and one tropical maize inbred (NC350) to sample the pool of inbreds comprising modern maize, for which high-quality genome sequences and annotations were available. We also selected the color converted W22 inbred, which is widely used for transposon mutagenesis, to identify transposon insertions in regulatory and coding regions. We also hoped to identify distal regulatory regions that might account for the very high proportion of SNPs that lie in intergenic regions found in Genome Wide Association Studies (GWAS)^9,24^, such as in the Nested Association Mapping population (NAM)^25^. B73 and NC350 are both NAM lines used in these studies. We also generated a high-quality reference genome assembly from the teosinte inbred line TIL11, using PacBio HiFi and BioNano Optical Mapping (Methods). A similar, but un-scaffolded assembly was published recently, revealing substantial intergenic transposon insertion variation between B73 and TIL11^26^. We subjected our high-quality assembly of TIL11 to the same annotation pipelines as the published maize inbred genomes for consistency (Extended data Fig. 1a). The TIL11 genome has several megabase-long inversions on chromosomes 1, 2, 4 and 7 conserved with all inbreds (Extended data Fig. 1b,c), likely representative of an event predating modern maize diversification. Furthermore, large differences between inbreds are also present, for example a duplication found only in W22 on chromosome 3, or a duplication found only in NC350 on chromosome 10 (Extended data Fig. 1c).

**Figure 1.**
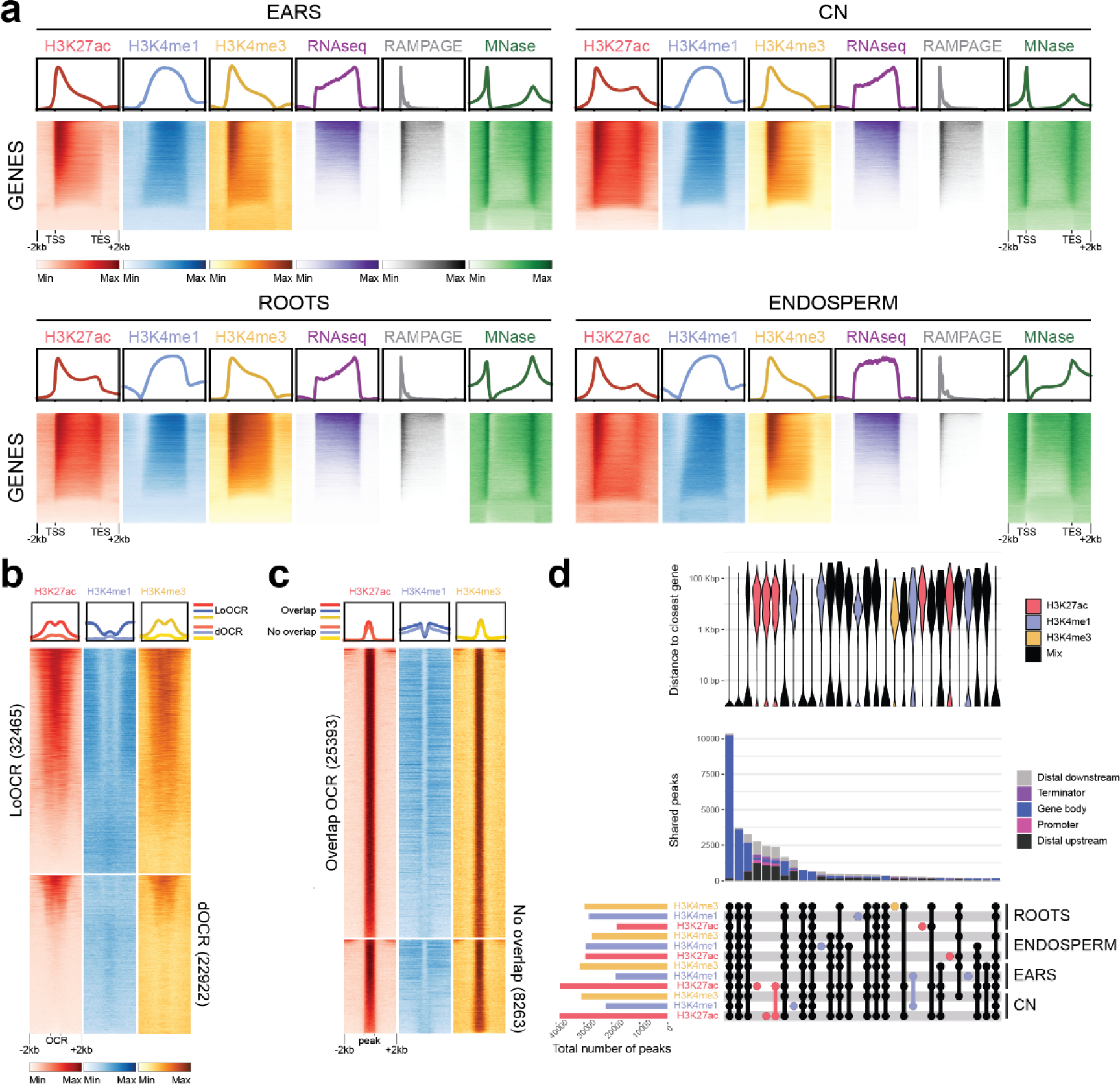
Histone H3 modifications mark DNA regulatory elements in maize inbred lines**. a.** Heatmaps and metaplots of H3K27ac, H3K4me1, H3K4me3, RNA-seq, RAMPAGE and MNase^27^ over all annotated genes in each tissue of B73 (NAM reference genome), scaled to the same size, with 2kb upstream and downstream. **b.** Heatmaps and metaplots of H3K27ac, H3K4me1 and H3K4me3 in local and distal open chromatin regions (LoOCR and dOCR, respectively) previously identified by ATAC-seq^30^. *Bona fide* regulatory elements are enriched for H3K27Ac and H3K4me3 but not H3K4me1. **c.** Heatmaps and metaplots of H3K27ac, H3K4me1 and H3K4me3 at all H3K27ac peaks (regulatory elements) in B73 ears. 25,393 peaks intersect previously identified OCRs but 8,263 peaks do not overlap, representing novel regulatory regions. **d.** Summary of shared CHIP-seq peaks in W22 (v2 reference genome). The Upset plot (lower panel) displays the overlap between H3K27ac, H3K4me1 and H3K4me3 peaks in the four tissues. The total number of peaks in each sample is shown on the histogram on the left-hand side of the intersection matrix, while the number of shared peaks between samples is shown above (middle panel), color coded by genomic feature. The violin plot (upper panel) compares the distance between peaks and the closest gene. Tissue specific peaks are mostly at distal elements, whereas loci with several histone marks in multiple tissues are mostly at annotated genes. Distal regulatory elements lie between 2kb and 100kb from the nearest gene.

MaizeCode data types conformed to ENCODE standards (Methods), but with plant-specific outcomes. Histone marks assessed by ChIP seq were H3K27ac and H3K4me3 (active enhancers and TSS), and H3K4me1 (gene bodies in plants) which complemented MNase accessibility data previously obtained from the same inbreds and tissues, as part of the MaizeCode project ^27^. RNA datasets included polyA+ RNAseq, RAMPAGE (5’ caps) and non-coding “short” RNA (or shRNA) of 150nt or less with 5’ tri- or mono-phosphate and 3’ hydroxyl groups. In plants, RAMPAGE and polyA+ RNAseq includes mRNA and ncRNA products of RNA polymerase II, while shRNA includes products of all 5 RNA polymerases, including 20-24nt siRNA (RNA Pol II and Pol IV), and longer transcripts generated by RNA Polymerase V^28,29^. RNA and chromatin samples were extracted from mature pollen, 5-10mm immature ears, 1-3mm root tips, endosperm harvested 15 days after pollination, and coleoptilar nodes 1 week after germination (Extended Data Table 1; Methods).

### H3K27ac marks genes and active enhancers bound by transcription factors

We identified active regulatory regions by integrating H3K4me1, H3K4me3 and H3K27ac ChIP-seq datasets with RNAseq, RAMPAGE and MNase accessibility datasets in up to all five tissues in the four inbreds (Extended data Table 1). The profiles of these histone marks over all genes (Fig. 1a) were similar to previously described patterns: peaks of H3K27ac, H3K4me3 and MNase at the TSS, and peaks of H3K4me1 over the gene body^9^ (Fig. 1a). These signals were present over transcribed genes, as shown by RAMPAGE and RNAseq signals in the different tissues tested (Fig. 1a) and in the different inbreds. We could confirm that these marks are enriched in OCRs previously identified by ATAC-seq in ears^30^ (Fig. 1b). The majority of local OCRs (LoOCRs, i.e. promoters and transcription start sites) were marked by H3K27ac and H3K4me3, while only a subset of distal OCRs (dOCRs) were marked by H3K27ac and H3K4me3, including those likely corresponding to the active cis-regulatory elements^9^ (Fig. 1b). Reciprocally, some H3K27ac peaks were not located in known OCRs, despite showing similar enrichment values in our experiments (Fig. 1c). Of note, whereas LoOCRs with high H3K27ac and H3K4me3 levels also showed H3K4me1 enrichment within 2kb up and downstream of the OCR, H3K4me1 was mostly absent from dOCRs, consistent with it exclusively marking gene bodies in plants (Fig. 1b).

We then investigated the tissue-specificity of these marks, by looking at the overlap between H3K27ac, H3K4me1 and H3K4me3 peaks in all tissues of the same inbred. Despite the variability in the number of peaks called in each sample, inherent to both the ChIPseq experiment and the peak calling algorithm, the largest sets of intersections corresponded to genic regions containing these 3 active marks in all four tissues investigated (Fig. 1d). This approach thus highlighted that most regulatory regions are shared between tissues, corresponding to the promoters of constitutively expressed genes. It also showed that when several marks were found together, they were mostly found in gene bodies, whereas when only one mark was present, notably H3K27ac, it was found in distal regions, which were located in a bimodal distribution centered around 2kb and 50kb upstream or downstream of the nearest gene (Fig. 1d).

In addition to histone modifications, we analyzed the binding profiles of selected transcription factors (TFs) in immature ears to illustrate how MaizeCode datasets can investigate mechanisms of domestication^8,10,11,31–33^. Almost all the binding sites (TFBS) coincided with H3K27ac peaks in at least one tissue (Fig. 2a). The majority of TFBS were within or close to a gene body, but about a third of the peaks overlapped distal enhancers (Fig. 2a). We also observed that each TF had a specific subset of targets, representing their unique regulatory networks (Fig. 2a). This observation was also highlighted by the fact that each TF had a preferential binding motif, which was representative of their family and DNA binding domain (Fig. 2b). The complex transcriptional regulation of development was emphasized by many enhancers that contained binding sites of multiple TFs, notably between FASCIATED EAR 4 (FEA4) and TU1-A. Interestingly, TFs often bound their own promoters and promoters of major domestication genes, while distal enhancers at the domestication genes *TB1*, *GT1*, *TGA1* and *RA1* were co-regulated by several TFs (Fig. 2c). These distal enhancers were also active in coleoptilar nodes, which include axillary buds, supporting previously identified branch suppression networks^12,34^. Overall, our analysis shows that H3K27ac peaks correlate well with active regulatory regions, whether marking TSS, proximal or distal enhancers, and harbor binding sites of developmental TFs whose functions have been refined during domestication.

**Figure 2.**
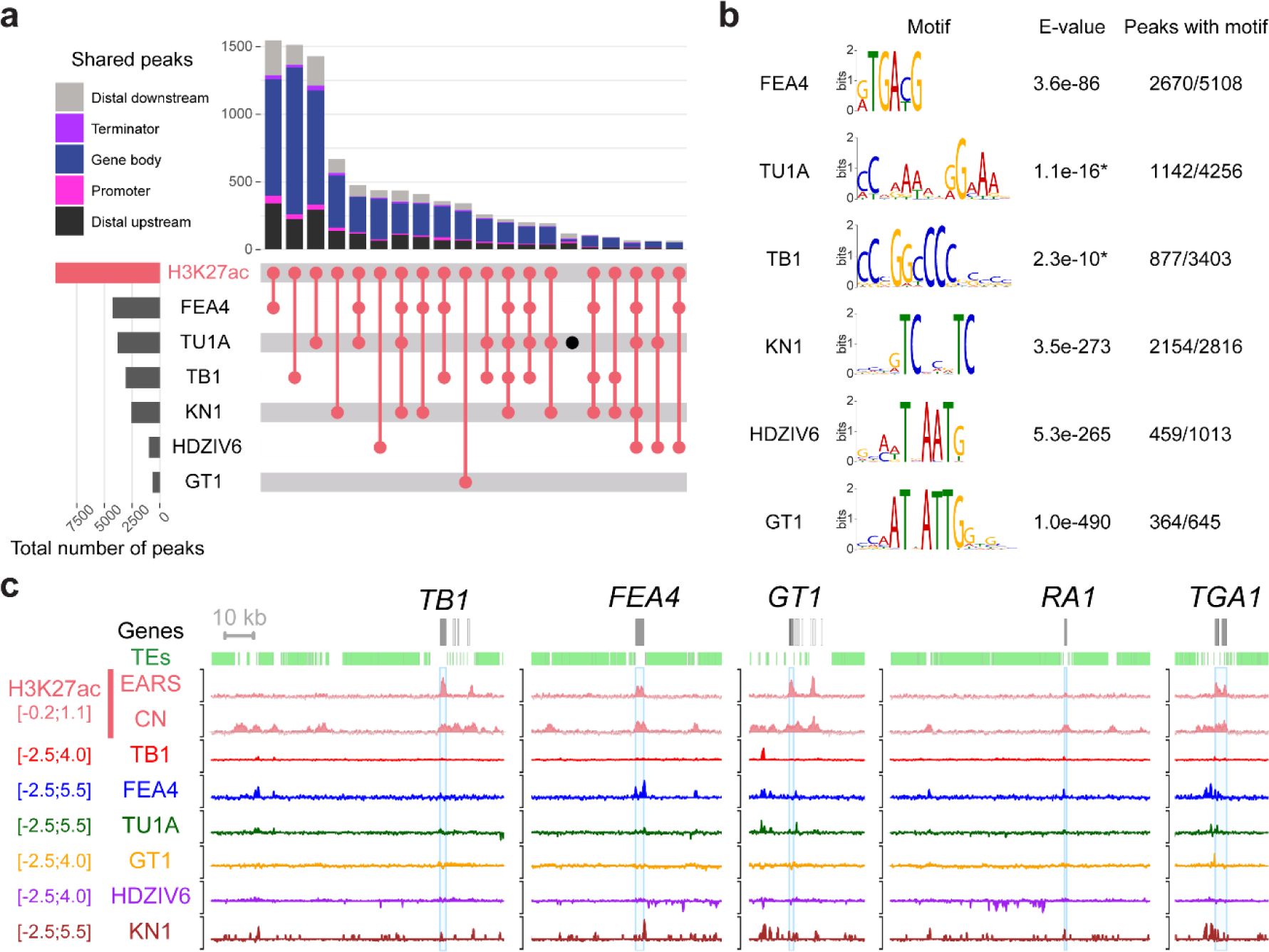
Enhancers at domestication genes are bound by transcription factor networks. **a.** Upset plot showing the overlap between H3K27ac peaks identified in B73 and binding sites of six transcription factors (TFBS) analyzed in this study. The total number of peaks called for each sample is shown on the histogram on the left-hand side. The number of shared peaks between the different samples are shown above the intersection matrix, each peak being colored by the genomic feature it intersects with. The majority of the TFBS are indeed within H3K27ac peaks, mostly overlapping gene bodies or at distal regions (>2kb from a gene) and highlights the interplay between these TFs. **b.** Best binding motif identified in each TF peaks with meme or streme (*)^76^. The motifs correspond to the respective family of each TF. **c**. Browser screenshots at major domestication loci (*TB1*, *GT1*, *RA1* and *TGA1*) as well as *FEA4*, which regulates a domestication trait, showing complex regulation of these developmental TFs with often co-regulation and auto-regulation.

### Pollen has a unique transcription profile for coding and non-coding regions

In parallel to the profiling of chromatin marks, we performed total RNAseq, RAMPAGE and short RNAseq in up to five tissues of the four inbreds (Extended data Table 1). The RNAseq data indicated that pollen had the most distinct gene expression profile, followed by endosperm (Fig. 3a; Extended data Fig. 2a-d). Thousands of genes were differentially expressed (DEGs) between each pair of tissues (Fig. 3a), with almost 4,000 and more than 5,000 genes consistently up and down-regulated, respectively, in pollen versus the four other tissues (Supplementary Information 1). By comparison, immature ears had only 260 and 191 genes up and down-regulated, respectively, versus all other tissues, whereas only 21 genes were down- regulated in the coleoptilar nodes compared to the four other tissues (Supplementary Information 1). This unique transcriptional profile of pollen was emphasized when clustering all the DEGs by their expression levels in each tissue, showing more extreme expression levels in pollen, consistently among all inbreds including TIL11 (Fig. 3a). Gene ontology (GO) analysis identified relevant DEGs in pollen, including genes involved in reproductive mechanisms in up-regulated genes (Fig. 3b). Similar GO terms were enriched in DEGs in other inbreds (Fig. 3b). Interestingly, genes involved in telomere maintenance were up-regulated in both pollen and in endosperm (Fig. 3d; Extended data Fig. 2e), the two tissues that engage in extended Breakage-Fusion-Bridge (BFB) cycles^35^, presumably due to aberrant telomere healing^36^. Interestingly, telomere maintenance genes were not upregulated in NC350 pollen or endosperm, suggesting regulatory variation potentially underlying variation in chromosome healing first described by McClintock^36^.

**Figure 3.**
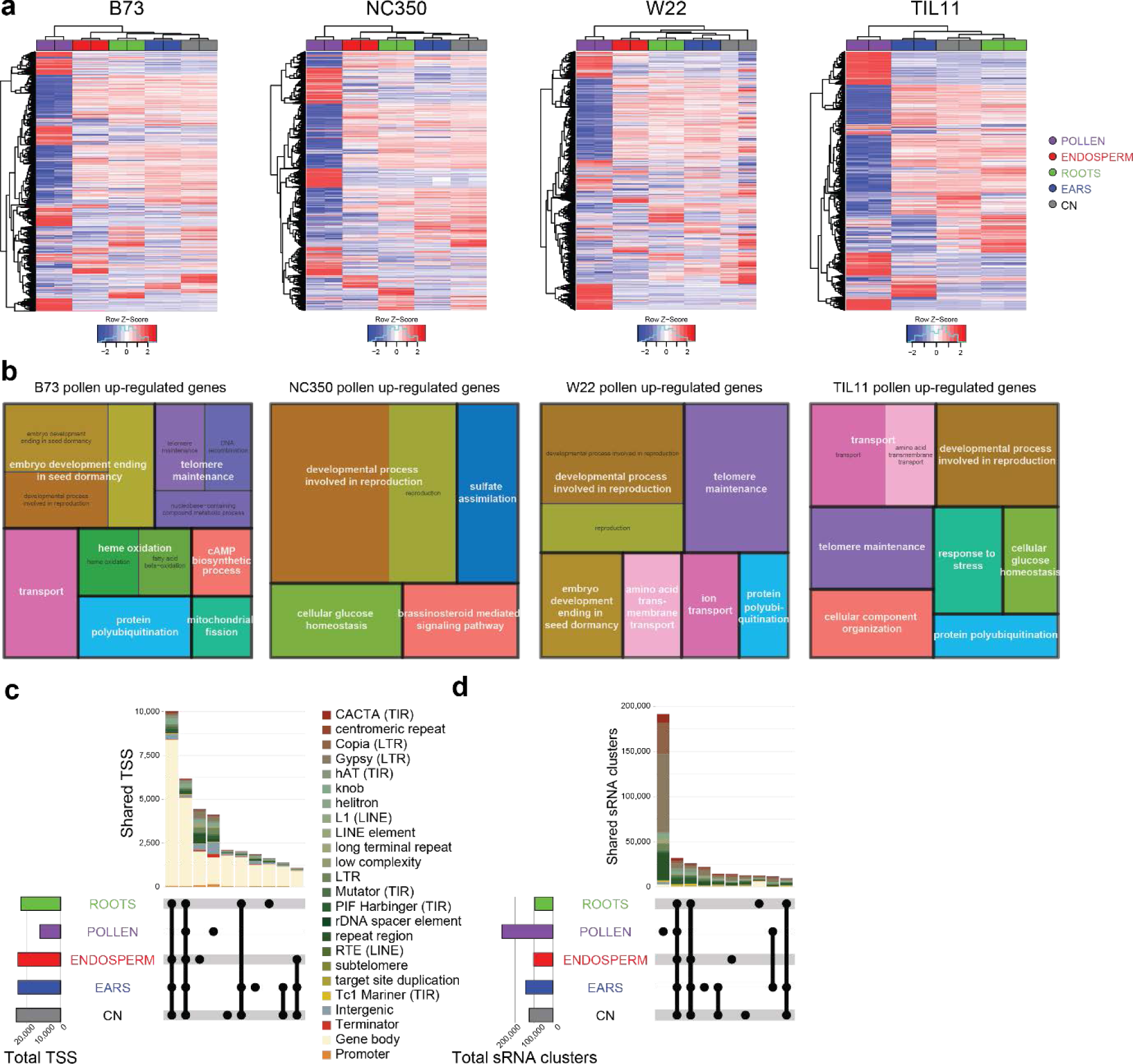
Pollen has a unique transcriptional profile compared to other tissues. **a.** Heatmap of all differentially expressed genes (DEGs) in each inbred and their expression level in each tissue (normalized z-score). **b.** Gene ontology (GO) terms enriched in genes up-- regulated in pollen versus all other tissues for each inbred. NC350 is missing genes involved in telomere maintenance. This difference is shared with endosperm (Extended Data Fig. 2) **c.** Upset plot of transcription start sites (TSS) identified by RAMPAGE in B73. The total number of TSS in each tissue is shown on the histogram on the left-hand side. The number of shared TSS between the different tissues are shown above the intersection matrix, color coded by genomic feature (including transposable element families). **e.** Upset plot of the sRNA clusters identified in shRNA-seq in B73. The total number of clusters in each tissue is shown on the histogram on the left-hand side. The number of shared sRNA clusters between the different tissues are shown above the intersection matrix, color coded by genomic feature (including transposable element families).

In addition to steady-state mRNA levels, we investigated the levels of capped RNA by RAMPAGE, which typically marks Transcription Start Sites (TSS)^37^. Consistent with RNAseq, pollen had the fewest TSS, about half of those shared between the other tissues (Fig. 3c). The large majority of loci (80%) shared between tissues mapped to annotated genes, but over 50% of the TSS unique to either pollen or endosperm were found in TEs and intergenic regions (Fig. 3c).

We next investigated levels of non-coding RNA by generating libraries of RNA molecules below 150nt in size with a 3’ hydroxyl and a 5’ phosphate that include canonical small interfering RNAs (siRNAs) generated by PolIV, as well as “short RNAs” (shRNA) longer than 30 nucleotides which included Pol V transcripts. The majority of siRNA clusters were unique to pollen and mapped almost exclusively to TEs, most notably long terminal repeat (LTR) retrotransposons (Fig. 3d). The second and third largest intersections were composed of clusters shared by all tissues, and shared by all tissues except pollen, respectively (Fig. 3d), further emphasizing the uniqueness of the pollen transcription profile, both coding and non-coding. Analysis of the size distributions of small RNAs revealed that B73 and TIL11 accumulated more 24nt than 21/22nt siRNAs in pollen (Extended Data Fig. 3), which were predominant in the other inbreds as previously reported for W22^38^. Intriguingly, immature ears of TIL11 had much lower levels of 24nt sRNAs, and higher levels of 21/22nt siRNA than found in maize, potentially due to differential activity of Dicer-like enzymes in teosinte^39^.

### Tissue and inbred-specific regulation of gene expression

The active marks studied here appeared to be reflective of transcription levels (Fig. 1a). To investigate this correlation, expressed genes were binned into 5 quintiles based on their expression levels (RPKM) to compare with enrichment levels of active histone marks (Extended data Fig. 4a). For all marks, a positive correlation could be seen, with highly expressed genes showing higher H3K27ac and H3K4me3 enrichment at the TSS, and higher H3K4me1 in the gene body. Interestingly, the differences between the Top 20% and the 20- 40% groups were not seen for H3K4me1, suggesting that a threshold was reached. Further confirming that these marks were associated with active genes in a tissue-specific manner, H3K27ac from immature ears was higher at genes up-regulated in ears versus all other tissues than at genes down-regulated in ears versus all other tissues (Extended data Fig. 4b). Conversely, the same genes showed the opposite pattern in other tissues, such as in roots where H3K27ac was higher in genes down-regulated in ears compared to other tissues (Extended data Fig. 4b). Similar trends were observed for H3K4me3, notably at the TSS, however H3K4me1 did not follow this trend. The same set of genes had higher H3K4me1 levels in both tissues, whether up-regulated in that tissue or not, suggesting that H3K4me1 was less variable across tissues and potentially not correlated with gene expression.

**Figure 4.**
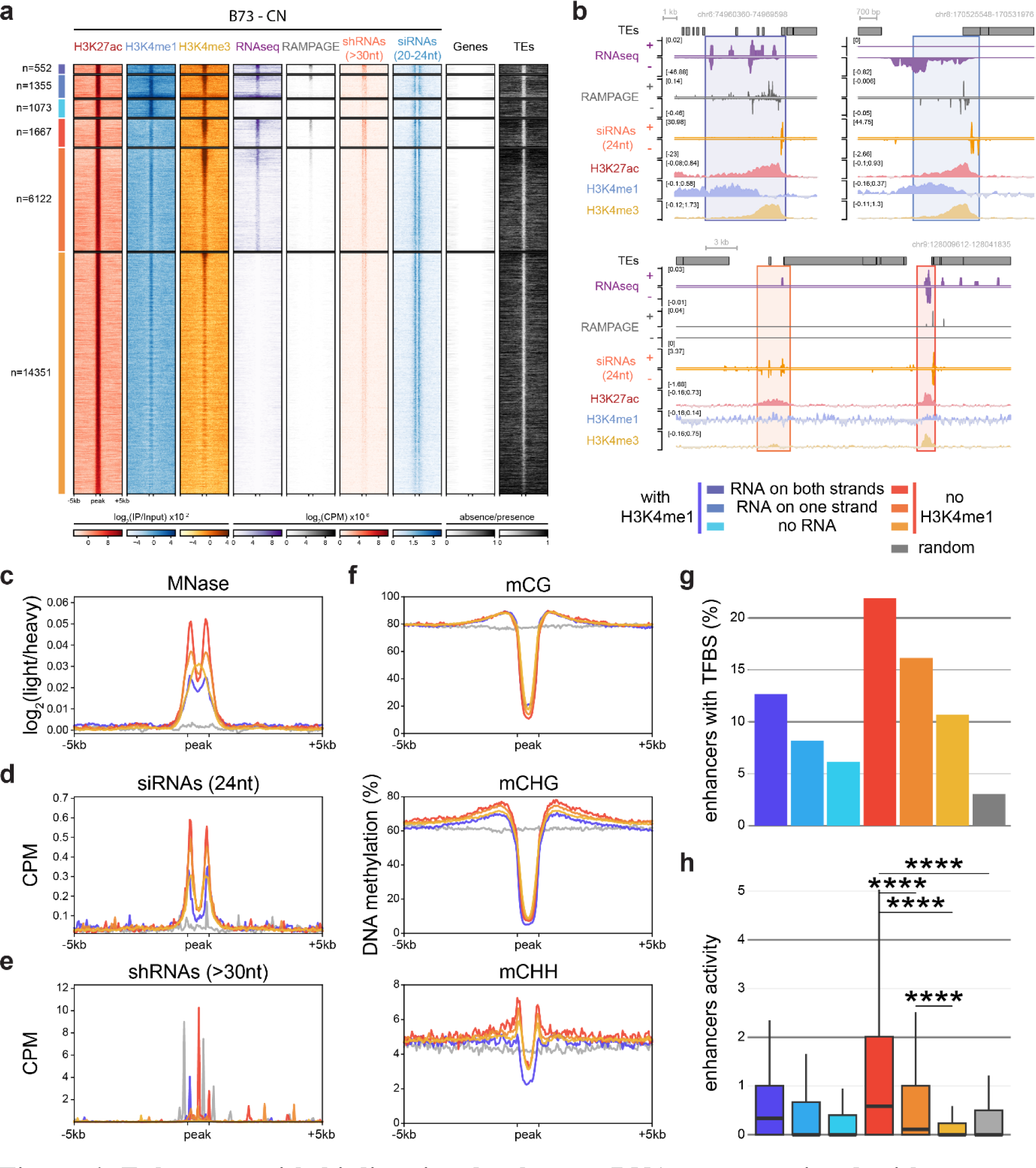
Enhancers with bi-directional enhancer RNAs are associated with stronger activity and higher RdDM at their boundaries. **a.** Heatmap of ChIP-seq and transcriptomic signals in B73 coleoptilar node (CN) at distal H3K27ac peaks and +/-5kb surrounding regions. Six classes of regulatory regions were identified based on the presence (blue) or the absence (red) of H3K4me1 peaks within 1kb, and on the presence of RNA-seq reads mapping to both strands, one strand, or none (from darker to lighter shades). The shRNA-seq datasets were split into longer fragments (>30nt) and canonical siRNAs (20-24 nt). Presence (black) and absence (white) of annotated genes and TEs surrounding the peaks are shown, demonstrating the absence of annotated features within regulatory regions. **b.** Browser screenshots of representative examples of uni- and bi- directionally expressed H3K27ac peaks (boxed), with (upper) and without (lower) H3K4me1 peaks. H3K4me1 peaks indicate the presence of unannotated genes. **c-f.** Metaplots at the three clusters without H3K4me1 peaks (red, as in **a**), the three clusters with H3K4me1 peaks merged together (blue), and random control regions (grey) of DNA accessibility (**c**) in micrococcal nuclease (MNase) from CN^27^, 24nt siRNAs (**d**) and short RNAs (>30nt) (**e**) generated in CN in this study, and DNA methylation in each sequence context (**f**) from seedlings^44^. These metaplots show that the bi-directional enhancers are more accessible regions with higher transcription levels of shRNAs, depleted of DNA methylation, but also more protected from neighboring TEs by targeting of RNA-directed DNA methylation by 24nt siRNAs. **g.** Percentage of peaks containing at least one transcription factor binding site (TFBS) from the TFs analyzed in this study. **h.** Measure of enhancer activity for each cluster by STARR-seq^9^. Bi-directionally expressed enhancers drive statistically higher transcription (maximum STARR-seq value within the enhancer) than uni-directional, not expressed or control regions (t-test, **** p<10^-5^).

In addition to tissue-specific expression, our data enabled comparison of tissue-specific expression in the different inbreds. *BOOSTER 1 (B1)* is a regulator of anthocyanin metabolism, and the *B1-I* allele engages in paramutation^40^. In addition to the hepta-repeat that is responsible for paramutation in *B1-I*, another tissue-specific enhancer is present about 45kb upstream from the TSS of the gene in B73^41^. This region was indeed marked by a H3K27ac peak in the coleoptilar nodes of B73 but not in the immature ears, correlating with *B1* expression and coleoptile pigmentation (Extended data Fig. 5a). In W22, the enhancer was slightly closer (∼20kb) to the transcription start site, due to structural variation caused by TEs, and the enhancer was active in immature ears, correlating with gene expression (Extended data Fig. 5b). This may be related to pigmentation of W22 under the control of *B1-bar*^42^.

**Figure 5.**
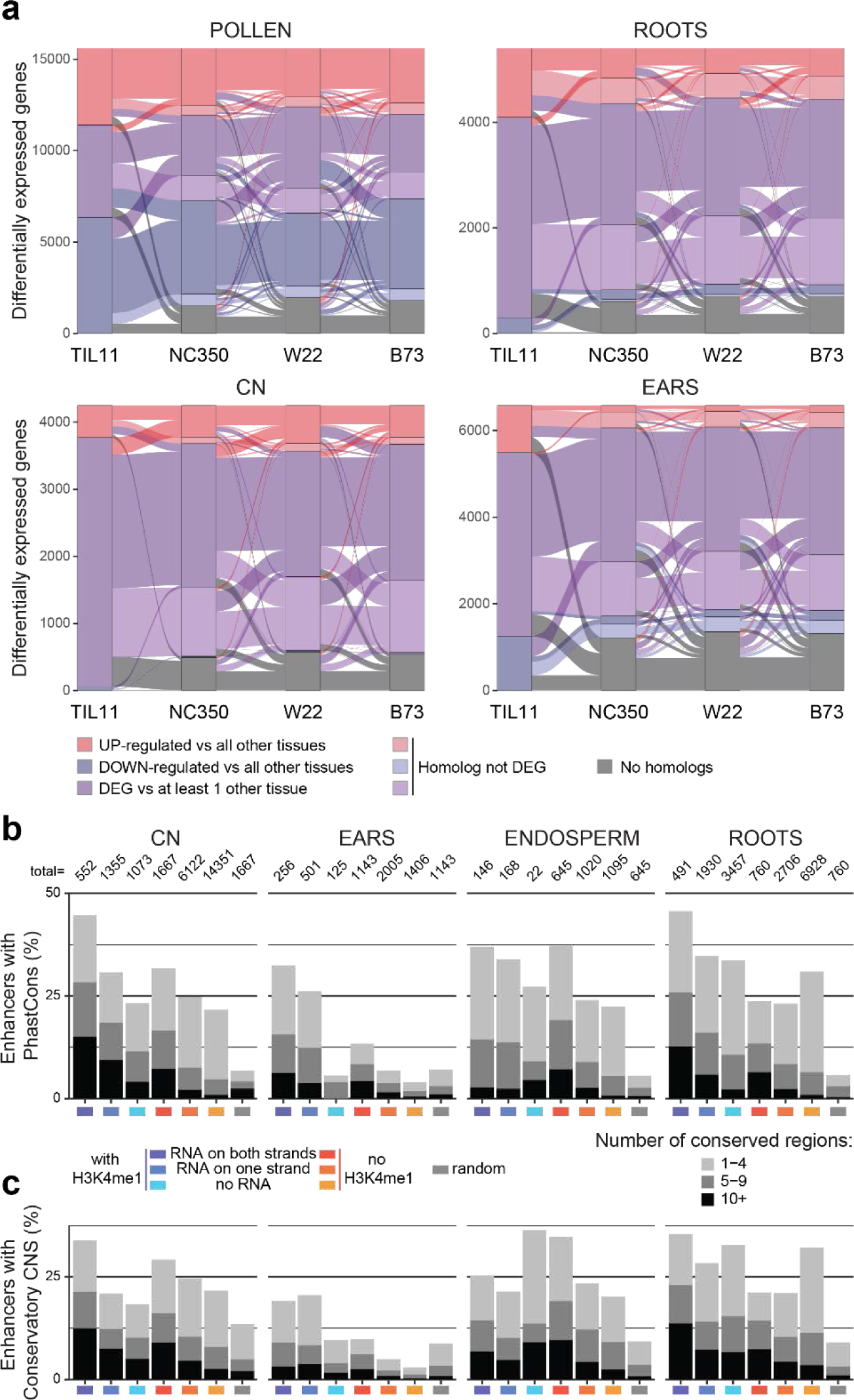
Domestication had a greater impact on transcription profiles and enhancer elements in ears. **a.** Alluvial plot showing the differentially expressed genes (DEGs) in four tissues of TIL11, and whether their homologs in modern maize maintain this differential expression. These plots show high level of transcription profile conservation in pollen, moderate levels in coleoptilar nodes (CN) and root tips, and low levels in immature ears, in addition to more genes not having a homolog (Methods). **b.** Percentage of enhancers containing conserved regions in the pan- andropoganeae clade identified by PhastCons (Methods). **c.** Percentage of enhancers containing conserved regions identified by Conservatory CNS (Methods). In both conservation analyses, misannotated genes show high levels of conservation as do the enhancers, especially the ones with bi-directional enhancer RNAs, in all tissues but in immature ears.

### Identification of enhancers with bi-directional enhancer RNAs

When focusing on the distal enhancers - as defined by H3K27ac peaks at least 2kb away from the closest annotated gene - we noticed that these distal enhancers with higher H3K27ac levels also had RNAseq and RAMPAGE signals. We expected some of these signatures to be caused by mis-annotations, either an unannotated gene or a wrongly annotated TSS. Since we noted that gene bodies were marked by H3K4me1, but that previously published distal OCRs are depleted in H3K4me1 (Fig. 1a,b), we intersected H3K27ac peaks with H3K4me1 peaks. We allowed the H3K4me1 peak to be within 1kb of the enhancers, to account for the distance between the TSS and the gene body (Fig. 4a; Extended data Fig. 6a). As expected, loci with both histone marks had the same molecular characteristics as genes (Fig. 4a,b), and were more often and more highly expressed in the corresponding tissue (Extended data Fig. 6a,b), thus likely representing misannotated genes.

Interestingly, many distal enhancers without H3K4me1 were also transcribed, either on one strand or on both strands (Fig. 4b; Extended data Fig. 6a). These bi-directionally expressed non-coding RNAs had additional molecular signatures reminiscent of animal enhancer RNAs (eRNAs), notably the presence of a 5’-cap, as shown by RAMPAGE (Fig. 4a,b; Extended data Fig. 6a,b). In addition, enhancers with bi-directional eRNAs had higher levels of H3K27ac and H3K4me3 (Extended data Fig. 6b), and corresponded to more accessible regions (Fig. 4c), shown by MNase accessibility data from the same tissues^27^. In maize, RNA-directed DNA methylation (RdDM) targets mCHH islands neighboring genes and cis-regulatory elements^43,44^. Consistently, we observed high levels of 24nt siRNAs at the borders of enhancers, especially those with bi-directional eRNAs (Fig. 4d; Extended data Fig. 6b), which accompany higher DNA methylation (Fig. 4f) in seedlings^44^. On the other hand, shRNAs (30- 150 nt), including presumptive Pol V transcripts, were mostly produced within the enhancer region (Fig. 4e; Extended data Fig. 6b).

Validating the importance of these enhancers in gene regulation, TFBS were more often in regions of bi-directional eRNAs than in control regions of similar sizes (Methods), or than in the misannotated genes with H3K4me1 (Fig. 4g). Furthermore, by comparing to previously published STARR-seq data^9^, we found that enhancers with bi-directional eRNAs had higher promoter activity than other distal enhancers or control regions (Fig. 4h). Despite their strong promoter activity, these enhancers appeared to be expressed mostly in a tissue-specific manner (Extended data Fig. 6c). These enhancers were also longer than enhancers without transcripts, but limited in size to several kilobases in all maize tissues, much shorter than “super-enhancers” in mammals which can be megabases long (Extended data Fig. 7a). The enhancer length did not seem to influence enhancer activity, since activity was higher in enhancers with bi- directional eRNAs in all tissues, whether measured by the maximum, the mean, or the median STARR-seq value in each enhancer (Extended data Fig. 7b). Further supporting the importance of these enhancers, enhancers with bi-directional eRNAs showed the highest overlap with OCRs identified by ATAC-seq in comparable immature ears from other studies^9,30^, often including 2 OCRs within the same enhancer (Extended Data Fig. 8a).

We then attempted to link these enhancers to the genes they regulate by intersecting with chromatin loops previously identified by chromatin conformation capture (Hi-C)^30^. Around half of enhancers with bi-directional eRNAs were present in chromatin loops, in similar proportions as previously identified distal OCRs (Extended Data Fig. 8b). Conversely, misannotated genes, marked by H3K4me1, were more often chromatin loop anchors (Extended Data Fig. 8b), which are enriched in gene-gene contacts. By comparison, local H3K27ac peaks were highly represented in gene-gene chromatin loops, slightly more (65% vs 60%) than local OCRs identified by ATAC-seq. Genes linked to enhancers with bi-directional eRNAs were more highly expressed than random genes, but only marginally more highly expressed than random genes present in chromatin loops (Extended Data Fig. 8c), which already represent a subset of highly expressed genes.

### Evolution of gene regulation during evolution and domestication

We also performed H3K27ac ChIP-seq in TIL11, and identified enhancers in immature ears and coleoptilar nodes (Extended data Table 1; Extended data Fig. 8). As expected, we found similar small RNA signatures in both TIL11 tissues, with 24 siRNAs targeting RdDM at the boundaries of active distal regulatory regions (Extended data Fig. 9).

We then set about comparing gene regulation between TIL11 and modern maize inbreds. Genes that were differentially expressed in each TIL11 tissue were compared to their closest homologs in other inbreds (Methods) to ask if they were also differentially expressed (Fig. 5a). As noted earlier, pollen had the most differentially expressed genes, and this pattern was observed in all inbreds including TIL11 (Fig. 3; Extended data Fig. 2). From the 10,531 DEGs in TIL11 pollen versus all other tissues, almost two thirds of their homologs were also DEGs in NC350 and B73 pollen versus all other tissues (64%, 62% respectively), and 54% were also DEGs in W22 pollen versus all other tissues (Fig. 5a). In coleoptilar nodes and in root tips, these proportions were slightly reduced, to about 40% and 30% respectively in the three inbreds. This proportion was drastically decreased in immature ears, where 10% or less of the genes differentially expressed in teosinte retained tissue-specificity in modern maize inbreds, including many novel genes in maize with no close homolog in teosinte (Fig. 5a; Methods). Overall, these results demonstrate that among the tissues studied here, tissue-specific gene expression evolved most rapidly in immature ears.

We next assessed the impact of domestication on tissue-specific cis-regulation by intersecting the different clusters of distal H3K27ac peaks identified above with conserved regions defined using PhastCons^45^ on the whole pan-andropogoneae clade (Methods). Between 25% and 50% of distal H3K27ac peaks neighboring H3K4me1 peaks had at least one - and often more than 10 - conserved regions, in all tissues, consistent with misannotated genes (Fig. 5b). The remaining enhancers also had a higher number of conserved regions, correlating with an increase in eRNA transcription (Fig. 5b). However, in ears, a much lower number of enhancers contained conserved regions, barely higher than the control regions (Fig. 5b). These results indicate that cis-regulatory elements driving tissue-specific expression in ears were impacted by domestication. To further examine the conservation of these enhancers, we used the conserved non-coding sequences (CNS) identified by the Conservatory Project^46^. Higher numbers of conserved regions were again found in enhancers with bi-directional eRNAs from all tissues, except from immature ears (Fig. 5c). This analysis supports the idea that these “super-enhancers” have been conserved throughout the Poaceae, but that the ones driving expression in the ears of modern maize are not conserved.

## Discussion

The MaizeCode initiative follows in the footsteps of the ENCODE project^15,16,47^ in cataloging regulatory regions in different tissues and inbred lines, so as to better understand the diversity of transcriptional regulation in maize. Additional datasets from teosinte enabled the analysis of tissue-specific transcription regulation in the context of domestication. In each inbred, most active histone marks were shared between all tissues tested (Fig. 1d). B73 immature ears had around 33,000 H3K27ac peaks in total (Fig. 1c), whereas more than 25,000 H3K27ac peaks were distal in coleoptilar nodes (Fig. 4a). It is likely that variation in the number of active enhancers in each tissue is caused by the heterogeneous composition of cell-types, a prediction borne out by single cell ATAC-seq studies of open chromatin regions (OCRs) in similar tissues^22^. Importantly, up to one quarter of the enhancers identified here in coleoptilar node by chromatin marks did not overlap with OCRs, consistent with a more comprehensive coverage of enhancers with histone marks^23^. In W22, however, immature ears and coleoptilar nodes had about 40,000 H3K27ac peaks, among which about 2,500 peaks were specific to each tissue, and about 2,500 were shared only between these two tissues (Fig. 1d). Despite such large variation, the molecular signatures at enhancers, and notably the presence of capped, poly-A tailed, bi-directional enhancer RNAs were identified in all inbreds and tissues studied here (Fig. 4; Extended data Fig. 6). The overall number of such “super-enhancers” was less variable between tissues (Extended data Fig. 6c), although we cannot exclude the possibility that enhancers more highly enriched in H3K27ac (Extended Fig. 6b) were the easiest to identify. It is important to note that variation in the quality of chromatin preparations inherent to the different tissues, with additional contributions from sequencing depth and the peak calling algorithm could have a large impact on the number of peaks, and thus enhancers, identified. Nonetheless, biological variation is very significant. For example, *BOOSTER 1* (*B1*) has a conserved regulatory region which is active in coleoptilar nodes in B73, but drives expression in immature ears in W22 (Extended data Fig. 5). *B1* is responsible for coleoptile pigmentation in B73, and glume pigmentation in tassels of W22. Differences in TE insertions in the region between the enhancer and the TSS in B73 and W22 could be responsible for this effect. Linking distal enhancers to their target genes can be done using chromatin conformation capture (Hi-C)^30^. We found that our enhancers were as often included in chromatin loops as OCRs marked by ATAC-seq (Extended Data Fig. 8b), yet the expression level of the genes they are contacting were only marginally increased compared to random genes forming loops (Extended Data Fig. 8c). Further studies would be required to more precisely associate enhancers to the genes they regulate, and to allow comparison between the different tissues and inbreds.

Transposable elements are drivers of cis-regulatory elements in plant genomes^48^, and the regulation of *tb1* is a well-known example of their impact on maize domestication^9,49^. TEs are under tight epigenetic control during the life-cycle of plants, and mechanisms responsible for keeping them in check include small RNAs and RNA-directed DNA methylation (RdDM)^50–52^. In maize, epigenetic signatures of mCHH methylation and 24nt siRNAs are found at gene boundaries, presumably to prevent euchromatic marks leaking into silenced TEs^43,44,53^. We also found that siRNAs and mCHH methylation are sharply elevated at the borders of distal regulatory elements in both modern maize and in teosinte (Fig. 4; Extended data Fig. 6,9). In addition, levels of siRNA, shRNA and mCHH were positively correlated with enhancer strength, being higher in enhancers with bi-directional eRNAs, than enhancers with eRNAs on one strand, than enhancers without eRNA (Fig. 4a,d; Extended data Fig. 6a,b). A possible explanation for this observation is that enhancer strength and increased DNA accessibility (Fig. 4; Extended data Fig. 6) enable easier access to RNA polymerase IV and Pol V (and therefore to the RdDM machinery) in the same way as they enable access to Pol II (Extended data Fig. 10). It is thus possible that the role of RdDM in maize is to prevent leakage of active transcription from enhancers into surrounding TEs with potentially detrimental impacts for the genome.

In mammalian genomes, H3K4me1 is associated with enhancers, whether poised or active^54,55^. In plants, H3K4me1 is associated with gene bodies^56,57^, and its pattern in all maize tissues and inbreds was consistent with previous studies^9^ (Fig. 1). In this study, we went one step further and used H3K4me1 as a proxy for genes (Fig. 4; Extended data Fig. 6). Validating this approach, distal H3K27ac peaks marked by H3K4me1 were more often present at chromatin loop anchors than *bona fide* enhancers, and than OCRs identified with ATAC-seq^30^ (Extended Data Fig. 8). This is consistent with previous work in maize that found that unmethylated regions of the genome (UMRs) with inaccessible chromatin had higher H3K4me1 levels than accessible ones, unlike H3K27ac and H3K4me3, which were higher in accessible UMRs^44^. H3K4me1 can be deposited in transcription-dependent and independent mechanisms^58^, potentially explaining the seemingly contradictory results of H3K4me1 being positively correlated with transcription, yet not being correlated with differential expression (Extended data Fig. 4). H4K16ac (as well as H2B ubiquitination) has been implicated in the recruitment of H3K4me1 methyltransferase^58^, and prevents chromatin remodeling by the epigenetic regulator DDM1^59^, which is found at RdDM targets in maize^60,61^. These observations suggest that H3K4me1 is present on genes that are not being silenced by DDM1, potentially allowing transcription elongation or preventing ectopic RdDM.

Consistent with the focus of breeding and domestication on yield and harvest traits, transcriptional regulation in immature maize ears showed very little conservation with teosinte, both in terms of patterns of expression of orthologous genes (Fig. 5a) and of their cis-regulatory elements (Fig. 5b,c). These results suggest that enhancers were not only reshuffled by TE insertions, as in the case of *tb1*, but evolved as rapidly as the genes they regulate, while maintaining their ability to drive strong transcription during domestication (Fig. 4e; Extended data Fig. 7b). The highest level of transcriptional conservation between maize and teosinte was found in pollen, despite having the most unique transcriptional profile of all the tissues examined for both coding and non-coding RNAs (Fig. 3; Extended data Fig. 2,3; Fig. 5a). This transcriptional profile is likely representative of conserved functions in reproduction, such as telomere maintenance, since breeding relies on fecundity and genome stability. It is also possible that conservation of pollen gene expression between maize and teosinte, which is otherwise unique among tissues, is the result of gene drive mechanisms such as *Teosinte Pollen Drive*^38^, that could be responsible for the fixation of epigenetic factors in modern maize varieties, and for the establishment of the molecular signatures identified here at their regulatory regions.

## Material and Methods

### Plant material and growth conditions

Seeds stocks for B73, W22 and NC350 were obtained from the Maize Genetics Stock Center, and for TIL11 from Dr. John Doebley.

For collecting immature ears, maize inbreds were grown in the CSHL Uplands Farm field in the summer until they reached the appropriate stage. The plants were collected from the field, and 5-10mm primary and secondary ear primordia were dissected in the lab then frozen in LN_2_ and stored at -80°C. TIL11 plants were grown in CSHL Upland Farm field from September to early October to promote floral transition by natural short day conditions. Immature TIL11 ears at an equivalent development stage (with inflorescence meristems, spikelet pair meristems, spikelet meristems and floral meristems) were collected under a dissecting microscope, frozen in LN_2_ and stored at -80°C.

For maize pollen samples, shedding tassels of field-grown plants as described above were bagged in the evening and mature pollen was collected the following day. After passing through a sieve to remove anthers, pollen was frozen in LN_2_ and stored at -80°C.

For harvesting TIL11 pollen, plants were grown in a short day (8h light/16h dark) walk-in chamber to promote floral transition. Fresh pollen was harvested, frozen in LN_2_ and stored at - 80°C.

For maize endosperm samples, ears of field-grown plants were sib-pollinated and collected 15 DAP. Endosperm was dissected in the lab, frozen in LN_2_ and stored in -80°C.

For maize and teosinte root tip samples, seeds were germinated on wet paper towels in a Pyrex dish in an incubator at 26°C in continuous darkness. After 5 days, 1-3 mm root tips were cut off with a razor blade on ice, frozen in LN_2_ and stored at -80°C.

For maize and teosinte coleoptilar nodes samples, seeds were germinated in flats in a long day (8h dark/16h light) growth chamber, 27°C day and 24°C night, and light at 130 μmoles. After 5 days, seedlings were unearthed and 5 mm sections around coleoptilar nodes were dissected on ice, frozen in LN_2_ and stored at -80°C.

At least three biological replicates of each tissue were collected.

### PacBio HiFi and ONT long-reads whole-genome sequencing of TIL11

Extracted DNA from TIL11 leaf nuclei was analyzed by Femto Pulse to assess fragment length distribution. For PacBio HiFi, DNA was sheared to ∼15kb using a Diagnode Megarupter following manufacturer’s recommendations. DNA was prepared for PacBio sequencing using the PacBio template prep kit 10. Briefly, 5ug of fragmented DNA prepared for sequencing via the PacBio kit, prepared libraries were size selected on Blue Pippin (Sage) from 10-15kb, and sequencing primer v2 was used. The library was loaded at 70pM on a PacBio Sequel II with a 48 hour movie. Circular Consensus processing was performed in SMRTLink to ensure multiple passes per fragment, and >=Q20 reads were selected for downstream assembly.

For ONT long reads sequencing, DNA was sheared to ∼30kb using a Diagnode Megarupter following manufacturer’s recommendations. DNA was prepared for Nanopore sequencing using the ONT 1D sequencing by ligation kit (SQK-LSK109). Briefly, 2ug of fragmented DNA was repaired with the NEB FFPE repair kit, followed by end repair and A-tailing with the NEB Ultra II end-prep kit. After an Ampure clean-up step, prepared fragments were ligated to ONT specific adapters via the NEB blunt/TA master mix kit. The library underwent a final clean-up and was loaded onto a PromethION PRO0002 flow cell per manufacturer’s instructions. The flowcells were sequenced with standard parameters for 3 days. Basecalling was performed with Guppy V5 to increase quality.

### Optical Map Generation

BioNano Optical Mapping was performed at Corteva Agriscience (Indianapolis, IN) following the protocols optimized for the NAM genomes^25^. Briefly, high molecular weight DNA was collected from fresh tissue from seedlings using the Bionano Prep™ Plant Tissue DNA Isolation Kit. Labeling was performed using the DLS Kit (Bionano Genomics Cat.80005) following manufacturer’s recommendations along with optimizations from the NAM samples. DNA was stained and quantified by adding Bionano DNA Stain to a final concentration of 1 microliter per 0.1 microgram of DNA. The labeled sample was then loaded onto a Bionano chip flow cell where molecules were separated, imaged, and digitized in the Saphyr System according to the manufacturer’s recommendations (https://bionanogenomics.com/support-page/saphyr-system/). Data visualization, processing, and DLS map assembly were conducted using the Bionano Genomics software Access, Solve and Tools.

### TIL11 Genome Assembly and Assessment

Prior to the genome assembly, we first assessed the size and heterozygosity of the TIL11 genome by analyzing the frequency distribution of 21-mers within the PacBio HiFi reads using KMC v3.1.1^62^ and GenomeScope v2.0^63^. This analysis confirmed the high quality of the HiFi reads and very low rates of residual heterozygosity (<0.001%) with on average 22x coverage in reads averaging 11.7kbp. Following this initial evaluation, we proceeded with the de novo assembly of long reads from PacBio HiFi Sequencing data using HiCanu^63,64^ optimized for high-fidelity long reads. The resultant assembly spanned 2.397 Gb with a contig N50 of 22.4Mbp (max: 95.0Mbp). These contigs were then scaffolded and packaged following the protocol used for the Maize NAM accessions^25^ using BioNano optical mapping data with the Bionano Access software and ALLMAPS. This yielded a highly contiguous & accurate, chromosome scale assembly with a scaffold N50 of 229.43Mbp and a contig N50 of 45.03Mbp. We assessed both consensus accuracy and completeness by analyzing the HiFi k-mer copy number spectra using Merqury version 2020-01-29^65^. Additionally, to gauge assembly completeness, we employed BUSCO v5.0.0^65,66^ with the embryophyta database from OrthoDBv10^67^ in genome mode. We investigated augmenting the assembly using the ONT long reads but found only potentially marginal improvements so did not include these results. Assembly based Structural Variants (SV) were characterized by aligning the chromosome scale assemblies of B73v5, NC350 and W22 lines to TIL11 using winnomap^68^ and further analyzing them using the SyRI package^69^.

### TIL11 annotations and gene orthology

Gene annotations for the TIL11 genome were done using the same protocol as described for the NAM genomes^25^. Orthologous genes were called using the Ensembl Compara Trees^70^. We dumped orthologs between two species from ensembl compara database with API. The orthology is a subclass of homology in the compara database. It was assigned by compara pipeline after reconciliation between gene tree and species tree. For any pair of homologs in a gene family, if their most recent common ancestor went through speciation event, these two homologs were deemed as orthologs. The annotation for TIL11 and comparative analysis with other NAM genomes is available on Gramene Maize (https://maize-pangenome.gramene.org/)

### Chromatin immuno-precipitation sequencing (ChIPseq) of histone modifications

The following amounts of tissues were used for each chromatin preparation: 10 coleoptilar nodes, 150 root tips, 10 immature ears and 10 endosperms. Chromatin was extracted as previously described^71^. Briefly, tissue was fixed in PBS with 1% formaldehyde under vacuum for 30 minutes. Crosslinking was stopped by adding glycine solution to 0.1 M final concentration. Fixed tissue was ground with pestle and mortar in LN_2_ and further disrupted using a dounce homogenizer. Chromatin was sheared using Covaris ultrasonicator and 300 μl of the chromatin prep was used for each immunoprecipitation with exception of coleoptilar nodes where 500 μl was used. The following antibodies were used to target chromatin modifications: H3K4me1 (Abcam, ab8895), H3K4me3 (Millipore, 07-473) and H3K27ac (Abcam, ab4729). Mixture of Dynabeads with proteins A and G (1:1) (Invitrogen) was used to pull-down the protein/DNA complexes and DNA was purified using ChIP Clean-up and Concentrator kit (Zymo Research). Libraries were constructed using Ultra II DNA kit (NEB).

### ChIP-seq of transcription factors

TU1-A-YFP, the dominant duplication ^11^, and GT1-YFP^10^ transgenic lines were introgressed into the *bd1;Tunicate* (*bd1;Tu*) double mutant background, which produces highly proliferative ears, to generate large amounts of ear tissue. ChIP experiments were adapted from a previously described protocol^32^. Briefly, two biological replicates of freshly harvested ear tissues were cross-linked in ice-cold buffer containing 10mM HEPES-NaOH PH7.4, 1% formaldehyde, 0.4 M sucrose, 1 mM EDTA, and 1 mM PMSF, for 20min under vacuum. Glycine was then added to a concentration of 0.1 M for another 5 min under vacuum to quench the crosslink. Nuclei extraction and immunoprecipitation were conducted as previously described^33^ using CelLytic PN Isolation/Extraction Kit (Sigma-Aldrich) and high-affinity GFP-Trap magnetic agarose (ChromoTek, gtma-20). ChIP-seq libraries were built as previously described^33^ using NEXTflex ChIP-seq Kit (PerkinElmer Applied Genomics) and AMPure XP beads (Beckman Coulter). ChIP-seq libraries were quantified by KAPA Library Quantification Kits (Roche) and sent for Illumina sequencing. ChIP-seq data generated from previous studies were used for ZmHDZIV6-YFP^33^, FEA4-YFP^32^, KN1^31^ and TB1^8^.

### Whole-transcriptome sequencing (RNA-seq)

For all inbreds and tissues, RNA was extracted with Direct-zol RNA Miniprep Kit (Zymo Research). 1 μg of total RNA was processed with the TruSeq Stranded Total RNA LT Kit (Illumina) as follows: all ribosomal RNAs were removed with RiboZero Plant included in the kit. After RNAclean XP purification (Beckman Coulter), anchored oligo(dT) and random probes were added and the RNA was fragmented. cDNA synthesis was performed followed by 2nd strand synthesis, 3’ adenylation, adapter ligation and target amplification. After purification with AMPure XP (Beckman Coulter) samples were quantified on a 2100 Bioanalyzer using a HS-DNA-Chip (Agilent), and adjusted to a concentration of 10 nM. Libraries were then pooled at equimolar concentration and sequenced on an Illumina NextSeq 500 Sequencer with a paired end 150 bp run.

### RNA Annotation and Mapping of Promoters for the Analysis of Gene Expression (RAMPAGE)

This protocol is a modified version of a previously published method^37^. Starting with 5 μg of total RNA, ribosomal RNAs were removed using the RiboMinus Plant Kit for RNA-Seq (Thermo Fisher Scientific) followed by incubation with Terminator 5′-Phosphate-Dependent Exonuclease (TEX) (Lucigen) to remove all residual RNAs containing 5’ monophosphate. We then performed first-strand synthesis using the SMARTer Stranded Total RNA Kit V2- Pico Input Mammalian (Takara). Following purification with RNAclean XP (Beckman Coulter), 5’ cap oxidation, 5’ cap biotinylation, RNase I digestion, and streptavidin pulldown (Cap Trapping) were performed as previously described^37^. Amplification of purified cDNAs (two rounds of PCR to attach Illumina adapters and amplify the libraries) followed by AMPure XP cleanup (Beckman Coulter) was done using the SMARTer Stranded Total RNA v2 kit (Takara), according to protocol. All samples were processed separately, quantitated on a 2100 Bioanalyzer using a HS-DNA-Chip (Agilent), and adjusted to a concentration of 10 nM. Libraries were then pooled at equimolar concentration and sequenced on an Illumina NextSeq 500 Sequencer.

### Total RNA Short RNA sequencing (shRNA-seq)

5 μg of total RNA were first depleted of rRNA with the RiboMinus Plant Kit for RNA-Seq (Thermo Fisher Scientific). RNA was de-capped using Cap-Clip Pyrophosphatase. The Illumina Truseq Small RNA protocol was used as follows: 3’ and 5’ adapters were ligated, followed by reverse transcription and amplification of the library. The BluePippin Size Selection system (Sage Science) was used to select library fragments ranging from 100 to 205 nt with the 3% agarose gel cassettes. Samples were quantified on a 2100 Bioanalyzer using a HS-DNA-Chip (Agilent), and adjusted to a concentration of 5 nM. Libraries were then pooled at equimolar concentration and sequenced on an Illumina NextSeq 500 Sequencer using a single end 150 bp run.

### Data analysis pipeline

Data analysis was performed using the Maizecode pipeline and accompanying scripts: https://github.com/martienssenlab/maize-code.

In brief, adapters were trimmed from raw sequencing files with cutadapt^72^ and data quality was assessed before and after trimming with FastQC. For ChIP-seq, trimmed files were mapped with bowtie2^73^ and processed with samtools^74^.

Peaks were called with Macs2^75^ and transcription factor motifs with the Meme suite^76^. For all samples, the two biological replicates were merged after mapping, split randomly into two pseudo-replicates and only the peaks called in the merged sample as well as in both pseudo- replicates were selected. For TB1, using the peaks identified in both biological replicates by Irreproducible Discovery Rate (IDR)^77^ generated more accurate results.

For RNA and RAMPAGE, trimmed files were mapped with STAR^78^. Differential gene expression analysis for RNAseq was performed with EdgeR v3.32.1^79^ and Gene Ontology analysis with rrvgo v1.5.3^80^ and topGO v2.42.0^81^. Transcription start sites were called with Macs2 using RNAseq as controls. For shRNA-seq, trimmed files were depleted of structural RNAs by mapping to rRNAs, snoRNAs and tRNAs with bowtie2. The unmapped reads were then mapped with Shortstack^82^. Mapped reads of 20 to 24nt were used to call sRNA loci with Shortstack, whereas reads longer than 30nt were kept for shRNA tracks. For DNA methylation, published datasets were processed with Bismark^83^. Browser tracks for all types of data were generated with Deeptools^84^. Heatmaps and metaplots were also generated with Deeptools, except for gene expression heatmaps which were produced with gplots v3.1.3; Upset plots were generated with ComplexUpset v1.3.3^85^; browser shots with Gviz v1.34.1^86^; boxplots with ggplot2 v3.4.1^87^. The following R packages and their versions were also used for data processing and plotting: dplyr v1.1.0; tidyr v1.3.0; cowplot v1.1.1; RColorBrewer v1.1.3; AnnotationForge 1.32.0; purrr 1.0.1; limma 3.46.0; stringr 1.5.0; wesanderson 0.3.6.

### Random control regions in mappable space

The B73 genome was fragmented into 150bp non-overlapping bins, which were then treated as single-end reads and mapped back to their respective genome following the same pipeline as ChIP-seq datasets. Only regions of the genome with at least one read mapped were kept as mappable. Bi-directionally expressed enhancers from each tissue individually were then randomly shuffled within this mappable space using the bedtools shuffle command^88^ in order to keep the same number and size distribution.

### Analysis of conservation within enhancers

An Andropogoneae phylogeny was inferred based on all genome-wide fourfold degenerate sites with <50% Androgoneae-wide missingness using RAxML. A neutral model of evolution was fit to this phylogeny using phyloFit from the PHAST 1.4 package^45^. A set of most conserved elements was generated using the PhastCons “most-conserved” flag from the PHAST package with an expected length of 40bp, after training to generate models of conserved and non-conserved elements using genome-wide multiple alignments with “--coverage 0.25”. To prevent reference-bias in the discovery of CNS, the B73 reference was masked and all other Tripsacineae were excluded for the phastCons analyses.

Conservatory CNSs were obtained from The Conservatory Project (www.conservatorycns.com)^46^.

## Acknowledgements

We thank Armin Scheben and Anat Hendelman for providing the files and expertise for the PhastCons analysis and the conservatory CNS analysis, respectively.

This work was funded by NSF IOS 1445025, as well as research in the Martienssen laboratory supported by the U.S. National Institutes of Health (NIH) grant R01 GM067014, the National Science Foundation Plant Genome Research Program, and the Howard Hughes Medical Institute and in the Jackson lab by NSF-IOS 2129189. The authors acknowledge assistance from the Cold Spring Harbor Laboratory Shared Resources, which are funded in part by a Cancer Center Support grant (5PP30CA045508).

## Authors contributions

JC, WRM, DW, DJ, MCS, TRG and RAM designed the study; MR, JL, XX, JD, MK and AS performed the experiments and the sequencing; JC, EE, CSA, SR, KC, SW and ZL analyzed the data and/or its significance; JC, DJ and RAM wrote the manuscript; MBH, WRM, DW, DJ, MCS, TRG and RAM acquired funding.

## Data and material availability

All sequencing datasets are publicly available on GEO SuperSeries: GSE254496.

## Code availability

All code is available on Github: https://github.com/martienssenlab/maize-code.

