## Extended Data Figures 1-10 for "MaizeCODE reveals bi-directionally expressed enhancers that harbor molecular signatures of maize domestication"

**a**

| Assembly Statistics TIL11 v1.1 | |
|---|---|
| **Contigs** | |
| # contigs | 9.803 |
| Longest (Mbp) | 109.55 |
| N50 (Mbp) | 45.03 |
| **Scaffolds** | |
| # scaffolds | 7,445 |
| > 10Mbp | 10 |
| Span (Mbp) | 2421 |
| Gapped bases (%) | 1.37 |
| **Annotation** | |
| # genes | 45,535 |
| Interspersed repeats (%) | 81.96 |
| **Quality Control** | |
| Merqury QV estimate | 66.9 |
| Merqury completeness estimate | 97.08 |
| Complete BUSCOs | 97.9 |
| Missing BUSCOs | 0.8 |

**b**

| TIL11 Structural variation summary | | | | |
|---|---|---|---|---|
| | | **B73** | **NC350** | **W22** |
| **Variant Type** | **Size** | | | |
| **Sniffles** | | | | |
| Deletion | 200bp - 1kb | 29,214 | 32,623 | 24,739 |
| | 1kb - 10kb | 46,191 | 45,333 | 36,448 |
| | > 10kb | 55,635 | 56,278 | 48,959 |
| Duplication | 200bp - 1kb | 578 | 553 | 1,052 |
| | 1kb - 10kb | 2,563 | 2,760 | 3,491 |
| | > 10kb | 14,783 | 15,155 | 13,481 |
| Inversion | 200bp - 1kb | 454 | 514 | 384 |
| | 1kb - 10kb | 1,352 | 1,407 | 1,272 |
| | > 10kb | 28,365 | 28,942 | 73 |
| Insertion | 200bp - 1kb | 22,183 | 25,078 | 21,061 |
| | 1kb - 10kb | 12,066 | 12,501 | 10,412 |
| | > 10kb | 174 | 205 | 178 |
| **Syri** | | | | |
| Inversion | > 1Mbp | 8 | 18 | 13 |

**c**

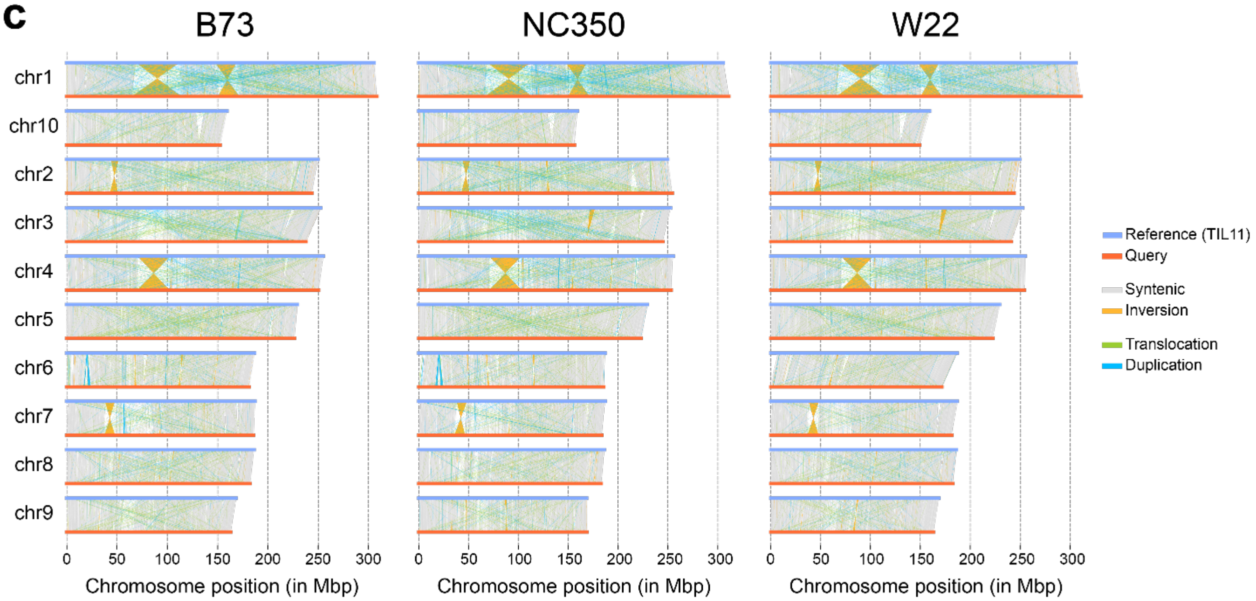


**Extended data Figure 1. TIL11 assembly statistics and structural variation with modern maize inbreds.**

**a.** Table of assembly and annotation statistics, showing high quality and comparable metrics with genomes assembled in the NAM project. **b.** Summary of the different structural variants between TIL11 and the three other inbreds studied. **c.** Full chromosome visualization of structural variants between TIL11 (used as reference) and the three other inbreds studied. Most large variants, and notably inversions, are conserved between maize inbreds.

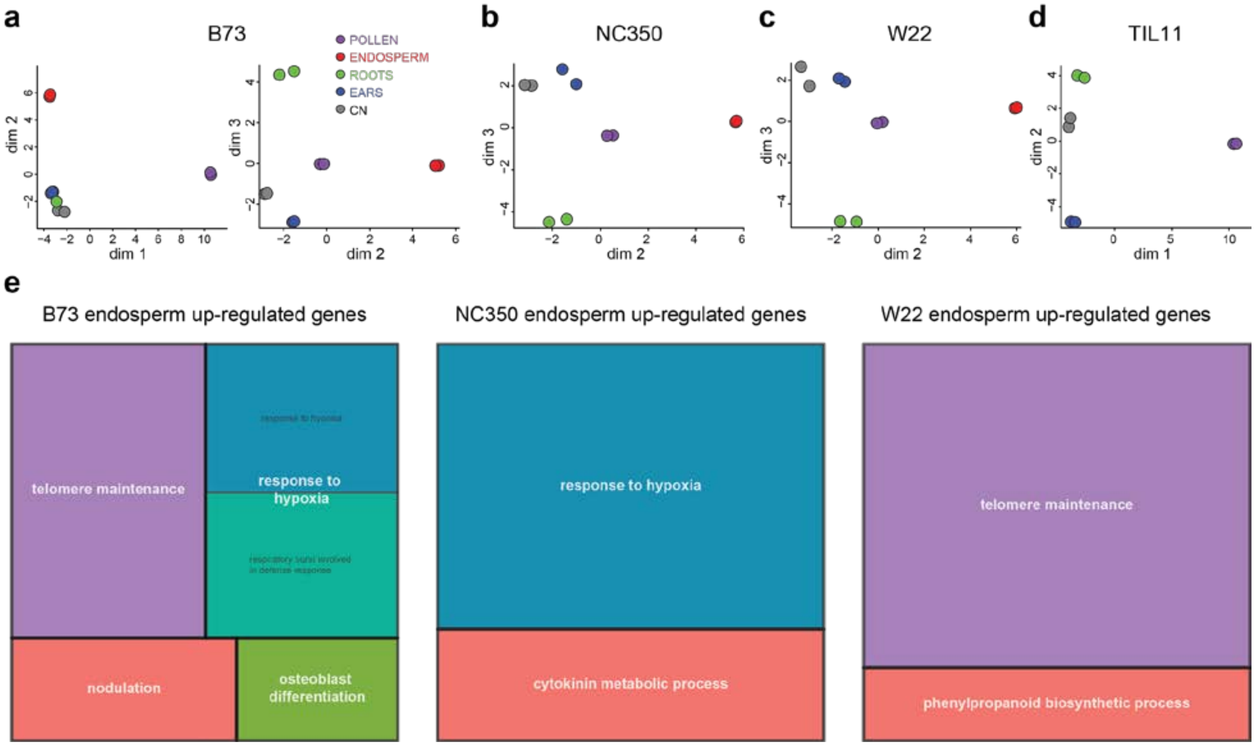


**Extended data Figure 2. Similar tissue-specific transcriptional profiles in maize and teosinte inbreds**.

**a-d.** Multidimensional scaling (MDS) plots of RNA-seq replicates in each of the 5 tissues of B73 (**a**), NC350 (**b**), W22 (**c**) and the 4 tissues of TIL11 (**d**). In all inbreds, biological replicates within samples cluster tightly together, while pollen and endosperm samples are most distinct. **e.** Gene ontology (GO) terms enriched in genes up-regulated in endosperm versus all other tissues of B73, NC350 and W22. Like pollen (Fig. 2), NC350 endosperm does not have an enrichment of genes involved in telomere maintenance.

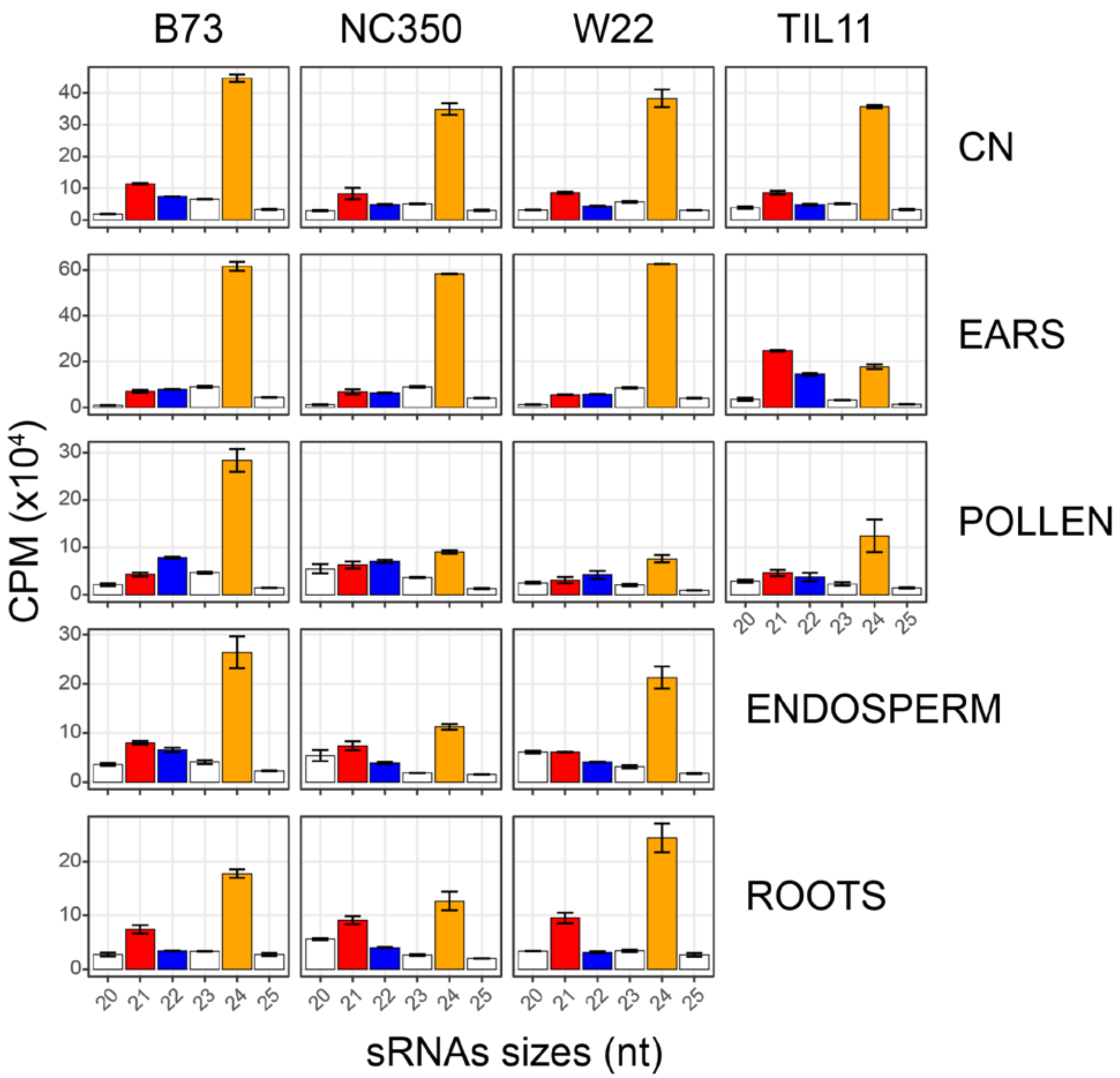


**Extended data Figure 3. Small RNA size distributions differ among tissues and inbreds.** Size distributions of sRNAs were calculated in each tissue of each inbred line (CPM, count per million mapped reads). Maize and teosinte inbreds have similar size distributions in coleoptilar nodes (CN), but differ in other tissues. In pollen, more 24nt sRNA (orange) accumulates in B73 relative to other inbreds, while in ears TIL11 has reduced levels of 24nt and increased levels of 21nt sRNAs (red). In root tips and endosperm, NC350 has reduced levels of 24nt siRNAs. Error bars are standard error between two biological replicates.

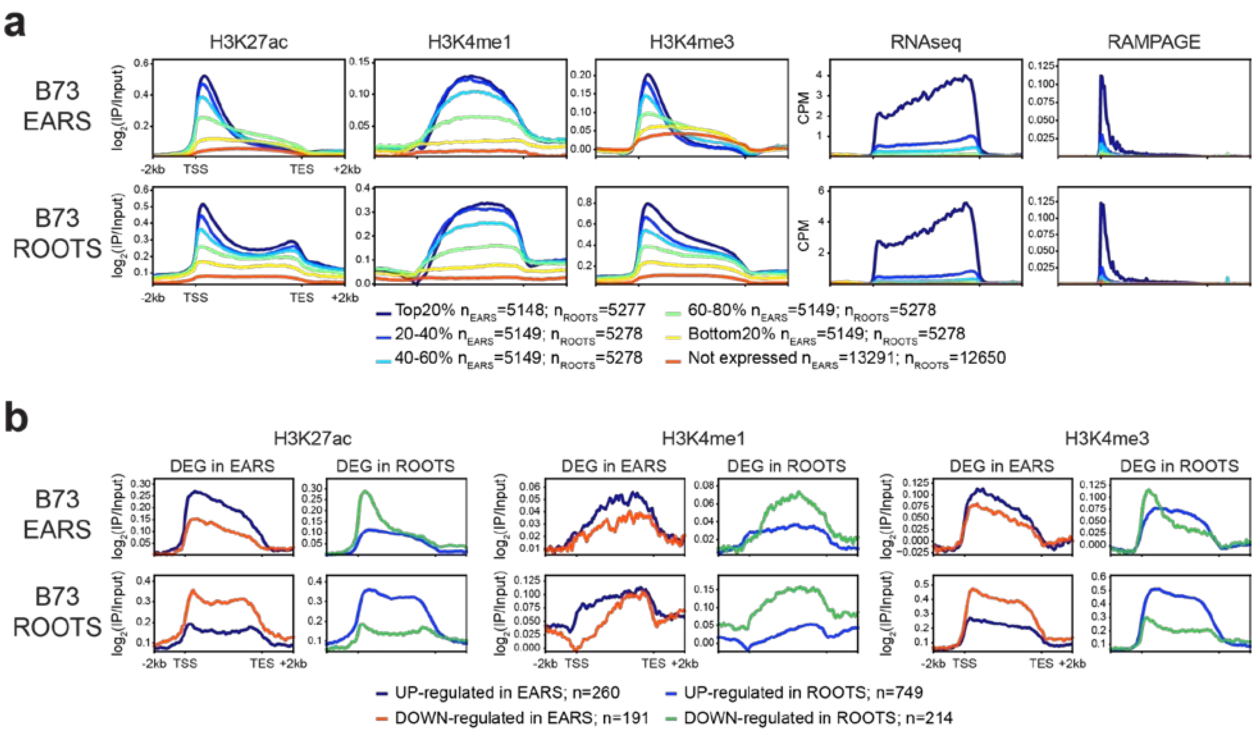


**Extended data Figure 4. Histone modification levels correlate with gene expression within tissues, but H3K4me1 does not correlate with differential expression between tissues.**
**a.** Metaplots of ChIP-seq, RNA-seq and RAMPAGE on genes grouped by expression levels within each tissue (RPKM). **b.** Metaplots of ChIP-seq in genes up or down-regulated in one tissue (immature ears or root tips) versus all other tissues.

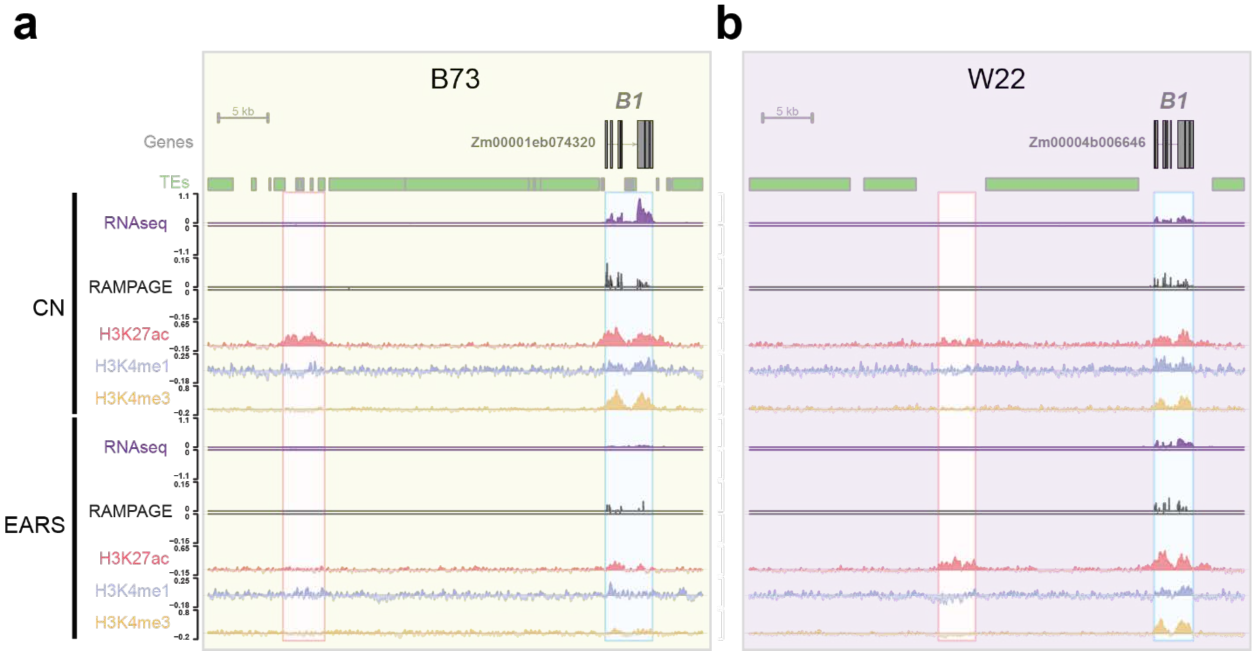


**Extended data Figure 5. Example of tissue-specific gene regulation in two inbreds: the *Booster1* (*B1*) locus.**

**a, b.** Browser shots of ChIP-seq ($log_2$[IP vs Input]) and transcriptomic data (CPM) in the *Booster 1* (*B1*) locus in coleoptilar nodes (CN) and immature ears of B73 (**a**) and W22 (**b**). In B73, the enhancer (red box) is active in the CN but inactive in immature ears, as shown by the peak of H3K27ac correlated with expression of the gene (blue box). In W22, the gene is expressed in both tissues, and the enhancer has H3K27ac signal in both tissues, which recapitulates the difference in pigmentation between the two lines.

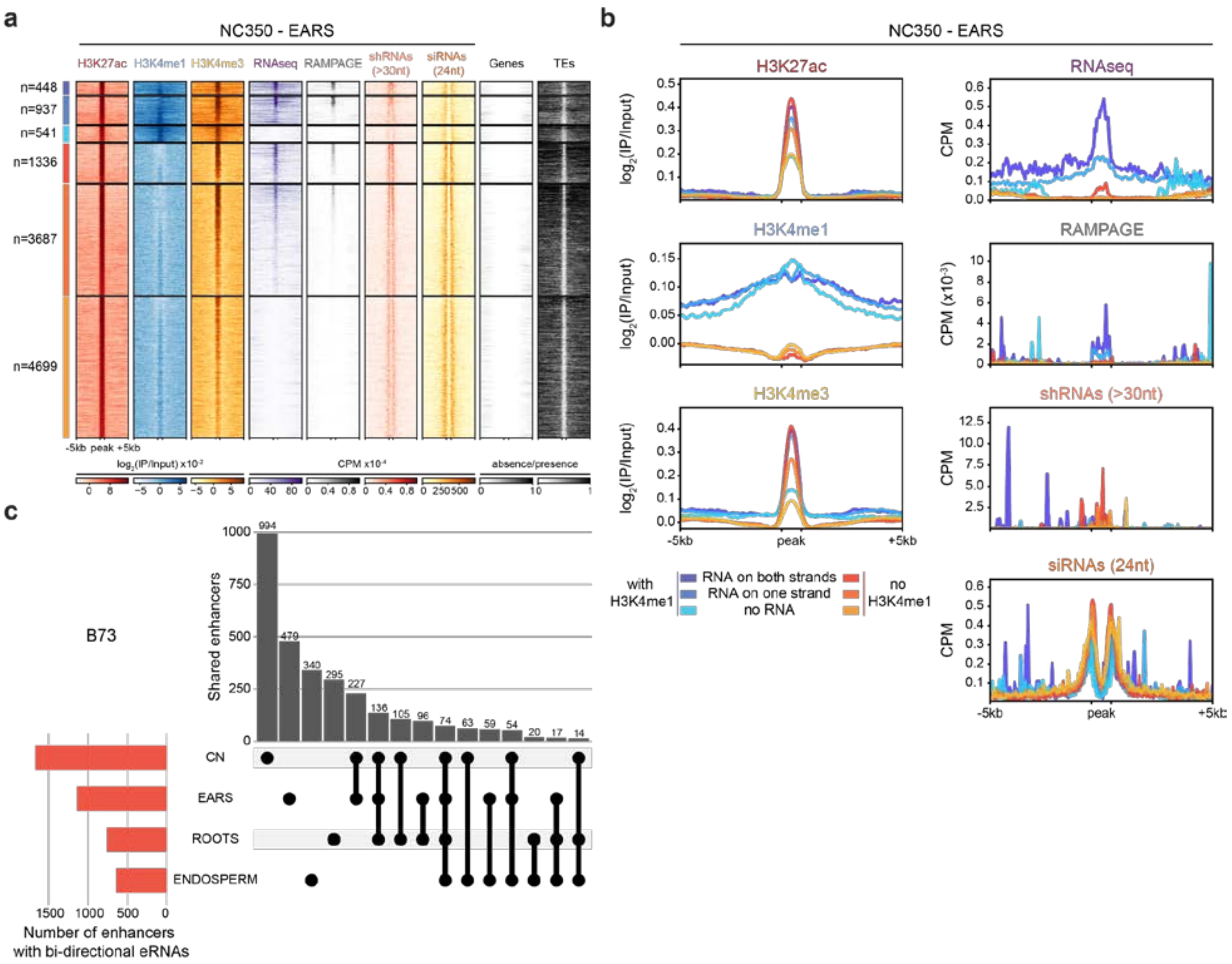


**Extended data Figure 6. Enhancers with bi-directional enhancer RNAs are tissue-specific.** **a.** Heatmap of ChIP-seq and transcriptomic datasets in NC350 immature ears at distal H3K27ac peaks. Six classes of regulatory regions were identified based on the presence (blue) or the absence (red) of H3K4me1 peaks within 1kb, and on the presence of RNA-seq reads mapping to both strands, one strand, or none (from darker to lighter shades). The shRNA-seq datasets were split into longer fragments (>30nt) labeled shRNA and canonical siRNAs (24nt).Presence (black) and absence (white) of annotated genes and TEs surrounding the peaks are shown, demonstrating the absence of annotated features within regulatory regions. **b.** Metaplots of the data summarizing the heatmap from (**a**). **c.** Upset plot of the enhancers with bi-directional enhancer RNAs identified in the four tissues of B73. The total number of enhancers identified in each tissue is shown on the histogram on the left-hand side. The number of shared loci between the different tissues are shown above the intersection matrix.

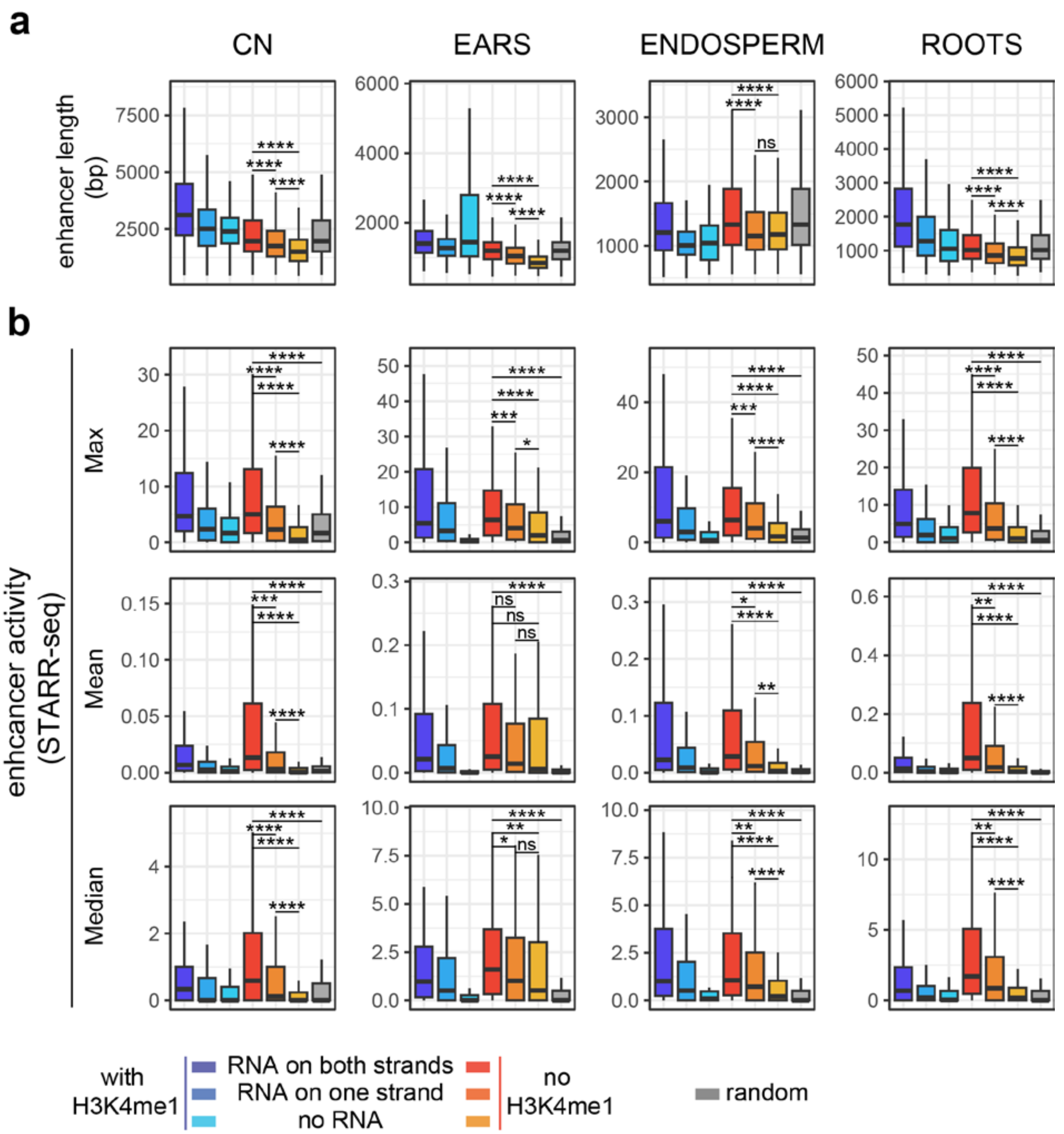


**Extended data Figure 7. Expressed enhancers are longer and inherently drive more expression, in all tissues.**

**a.** Size distribution of the distal H3K27ac peaks in each tissue of B73, split by the presence of H3K4me1 peak nearby, and the presence of RNA within the peaks. A set of control regions consisting of the bi-directional enhancers shuffled in the genome was included (t-test, * $p<10^{-2}$; ** $p<10^{-3}$; *** $p<10^{-4}$; **** $p<10^{-5}$; ns not significant). **b.** Distribution of enhancer activity of the same clusters than in (**a**), as measured by STARR-seq [9]. The maximum value found in each enhancer, the mean value of each enhancer, and the median value of each enhancer were plotted and always show higher activity of enhancers with bi-directional eRNAs (t-test, * $p<10^{-2}$; ** $p<10^{-3}$; *** $p<10^{-4}$; **** $p<10^{-5}$; ns not significant).

**a**

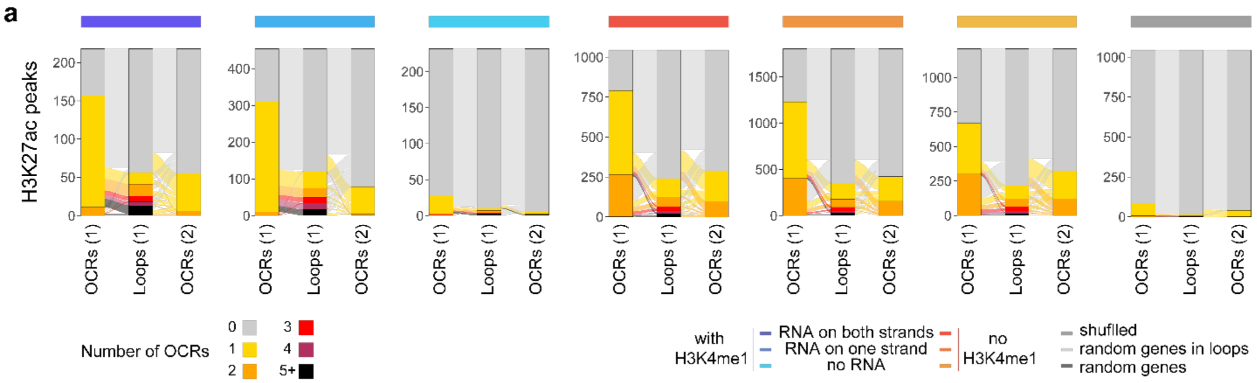


**b**

| | Total | Gene-gene (n=60,172) | Intergenic-gene (n=20,078) | Intergenic-intergenic (n=4,318) |
|---|---|---|---|---|
| Local OCRs (1) | 32,465 | 19,656 ( **60.55%** ) | 6,188 ( **19.06%** ) | 6 ( **0.02%** ) |
| Local H3K27ac | 25,218 | 16,602 ( **65.83%** ) | 5,218 ( **20.69%** ) | 48 ( **0.19%** ) |
| Distal OCRs (1) | 22,922 | 5,078 ( **22.15%** ) | 7,038 ( **30.70%** ) | 2,558 ( **11.16%** ) |
| bi-directional | 218 | 46 ( **21.10%** ) | 97 ( **44.50%** ) | 34 ( **15.60%** ) |
| uni-directional | 454 | 77 ( **16.96%** ) | 190 ( **41.85%** ) | 66 ( **14.54%** ) |
| no expression | 232 | 5 ( **2.16%** ) | 28 ( **12.07%** ) | 14 ( **6.03%** ) |
| bi-directional | 1,044 | 76 ( **7.28%** ) | 318 ( **30.46%** ) | 119 ( **11.40%** ) |
| uni-directional | 1,805 | 104 ( **5.76%** ) | 483 ( **26.76%** ) | 210 ( **11.63%** ) |
| no expression | 1,208 | 67 ( **5.55%** ) | 306 ( **25.33%** ) | 149 ( **12.33%** ) |
| shuffled | 1,044 | 156 ( **14.94%** ) | 89 ( **8.52%** ) | 22 ( **2.11%** ) |

**c**

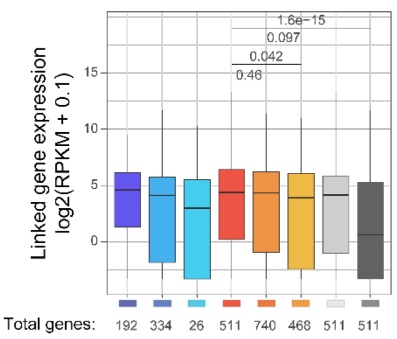


**Extended data Figure 8. H3K27ac-marked regulatory regions are present in chromatin loops to similar or higher levels than open chromatin regions defined by ATAC-seq.**

**a,** Alluvial plots showing the number of OCRs contained in H3K27ac peaks, split by the presence of H3K4me1 peak nearby, and the presence of RNA within the peaks. H3K27ac peaks identified in B73 immature ears were compared to open chromatin regions (OCRs) from ATAC-seq and to chromatin loops from Hi-C from (1) Sun et al. 2020[30] and to OCRs from (2) Ricci et al. 2019[9]. The highest overlap is between Sun et al. 2020 OCRs and enhancers with bi-directional enhancer RNAs (eRNAs). **b,** Table summarizing the number of enhancers found in the chromatin loop anchors identified by Hi-C[30]. H3K27ac peaks within 2kb of a gene body (local H3K27ac, green) is more often in a loop than local OCRs. Distal H3K27ac peaks are included in Intergenic loops to similar levels than OCRs. The presence of H3K4me1 however increases the percentage of these regions to be within loops, which support their classification as misannotated genes. **c,** Expression level in immature ears ($\log_2$(RPKM+0.1)) of the genes linked by chromatin loops to the different types of enhancers described in **a**. Genes linked to enhancers with bi-directional eRNAs are more highly expressed than random genes, but marginally more highly expressed than genes in loops (t-test).

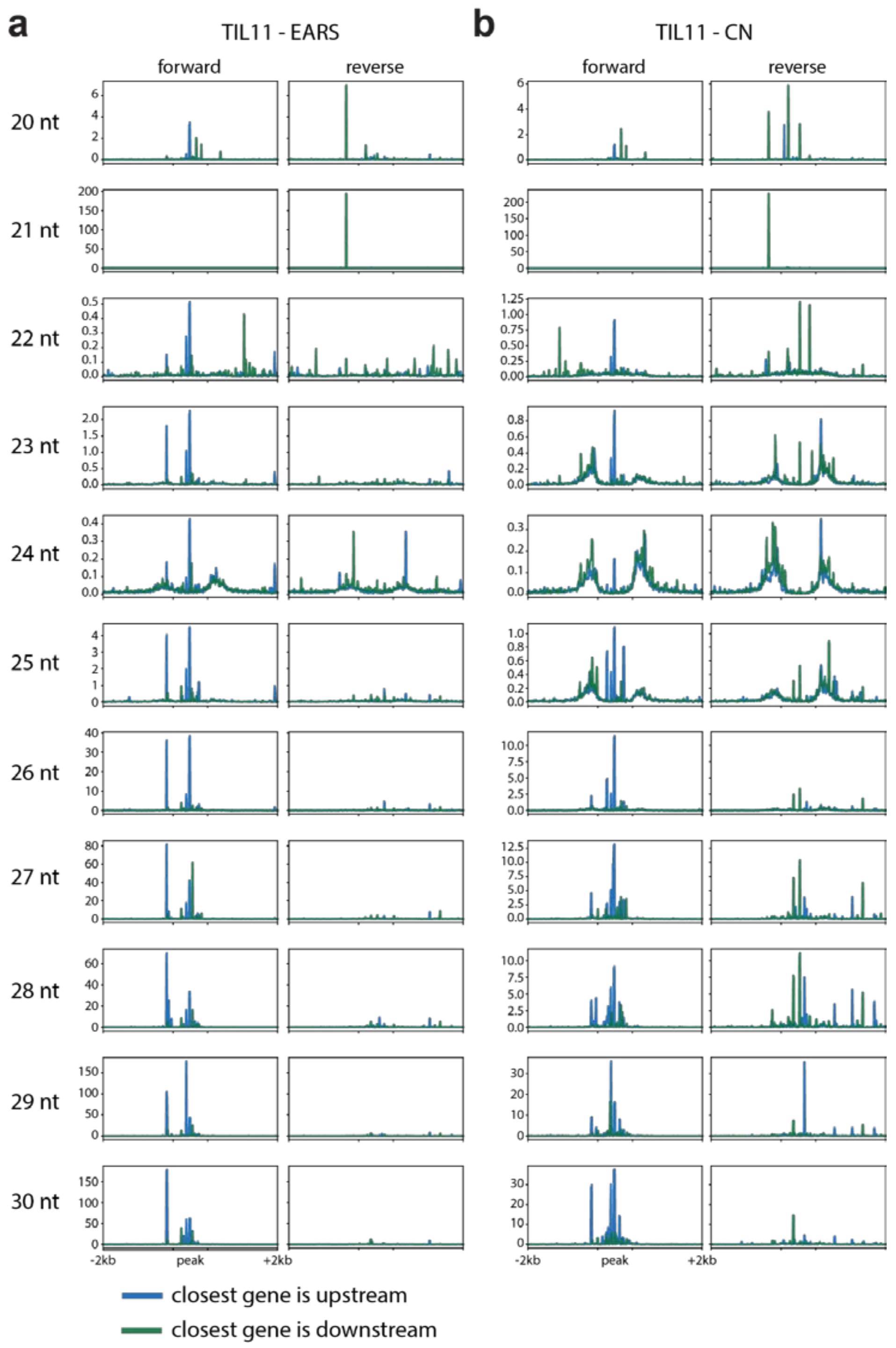


**Extended data Figure 9. Small interfering RNAs also target boundaries of enhancers in TIL11.**

**a,b.** Metaplots of short RNA data from TIL11 immature ears (**a**) and coleoptilar nodes (CN) (**b**) at distal enhancers identified in TIL11 immature ears. Small RNAs were split by read length from 20 to 30 nucleotides and mapping strands (forward or reverse) before normalization in count per million (CPM), and plotted on H3K27ac peaks which were further than 2kb from the closest gene, either upstream (green) or downstream (blue) from it.

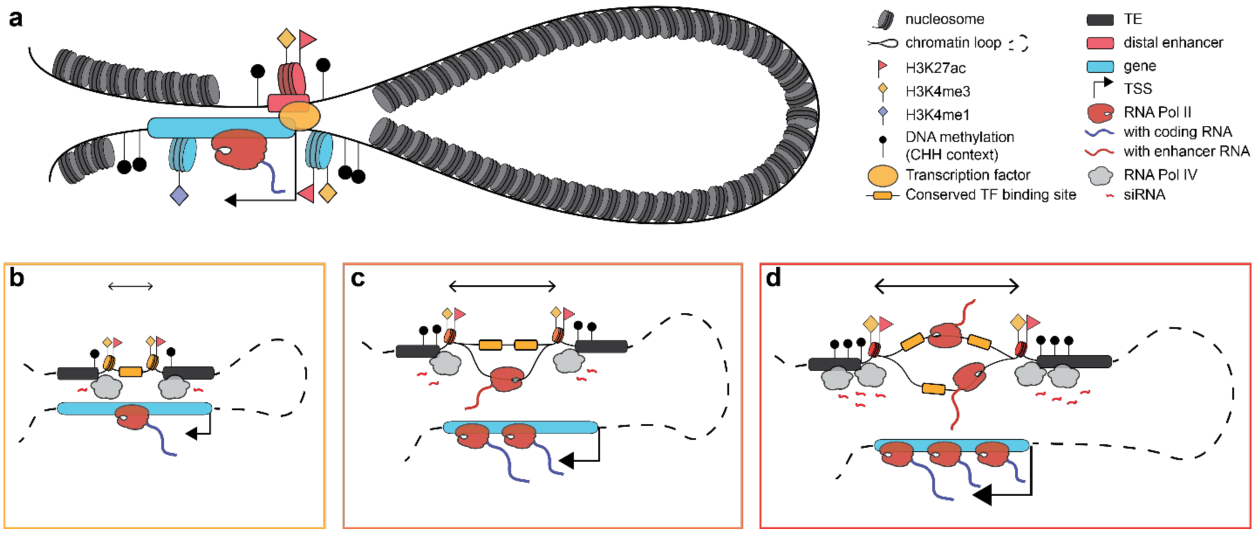


**Extended Data Figure 10. Model of enhancers with bi-directional enhancer RNAs.**

**a.** Distal enhancers, marked with H3K27ac and H3K4me3, form chromatin loops with the genes they regulate through transcription factors recruiting RNA polymerase. Genes are also marked by H3K27ac and H3K4me3 at their promoter, and contain H3K4me1 along the gene body. Both genetic features are surrounded by transposable elements and have CHH DNA methylation (mCHH) deposited by the RdDM machinery at their boundaries. Enhancers with higher number of transcription factor binding sites promote higher gene expression (shown by more RNA PolII proteins and bigger TSS arrows) which is correlated with larger, more accessible regions (bigger horizontal arrow) and an increase in transcription of enhancer RNAs from: **b**, no enhancer RNA (eRNA); **c**, eRNA on one strand; and **d**, bi-directional eRNAs. This increase in accessibility is visible at the enhancer with an elevated level of H3K27ac and H3K4me3 (larger symbols), and higher RdDM activity and mCHH at the boundaries (more Pol IV producing siRNAs and black lollipops), protecting transcription to extend into the neighboring transposable elements.
