## Extended Data Table 1 for "MaizeCODE reveals bi-directionally expressed enhancers that harbor molecular signatures of maize domestication"

**Extended data Table 1.** Datasets generated by the MaizeCODE consortium and analyzed in this study.

| | | RNA | | | ChIP-seq | | | TF ChIP-seq | | |
|---|---|---|---|---|---|---|---|---|---|---|
| **Inbred** | **Tissue** | **RNAseq** | **RAMPAGE** | **shRNAseq** | **H3K27ac** | **H3K4me3** | **H3K4me1** | **GT1** | **HDZIV6** | **TU1A** |
| B73 | Immature ears | 2 Reps | 2 Reps | 2 Reps | 2 Reps | 2 Reps | 2 Reps | 2 Reps | 2 Reps | 2 Reps |
| B73 | Endosperm | 2 Reps | 2 Reps | 2 Reps | 2 Reps | 2 Reps | 2 Reps | - | - | - |
| B73 | Root tips | 2 Reps | 2 Reps | 2 Reps | 2 Reps | 2 Reps | 2 Reps | - | - | - |
| B73 | Coleoptilar node | 2 Reps | 2 Reps | 2 Reps | 2 Reps | 2 Reps | 2 Reps | - | - | - |
| B73 | Pollen | 2 Reps | 2 Reps | 2 Reps | - | - | - | - | - | - |
| W22 | Immature ears | 2 Reps | 2 Reps | 2 Reps | 2 Reps | 2 Reps | 2 Reps | - | - | - |
| W22 | Endosperm | 2 Reps | 2 Reps | 2 Reps | 2 Reps | 2 Reps | 2 Reps | - | - | - |
| W22 | Root tips | 2 Reps | 2 Reps | 2 Reps | 2 Reps | 2 Reps | 2 Reps | - | - | - |
| W22 | Coleoptilar node | 2 Reps | 2 Reps | 2 Reps | 2 Reps | 2 Reps | 2 Reps | - | - | - |
| W22 | Pollen | 2 Reps | 2 Reps | 2 Reps | - | - | - | - | - | - |
| NC350 | Immature ears | 2 Reps | 2 Reps | 2 Reps | 2 Reps | 2 Reps | 2 Reps | - | - | - |
| NC350 | Endosperm | 2 Reps | 2 Reps | 2 Reps | 2 Reps | 2 Reps | 2 Reps | - | - | - |
| NC350 | Root tips | 2 Reps | 2 Reps | 2 Reps | 2 Reps | 2 Reps | 2 Reps | - | - | - |
| NC350 | Coleoptilar node | 2 Reps | 2 Reps | 2 Reps | 2 Reps | 2 Reps | 2 Reps | - | - | - |
| NC350 | Pollen | 2 Reps | 2 Reps | 2 Reps | - | - | - | - | - | - |
| TIL11 | Immature ears | 2 Reps | 2 Reps | 2 Reps | 2 Reps | - | - | - | - | - |
| TIL11 | Coleoptilar node | 2 Reps | 2 Reps | 2 Reps | - | - | - | - | - | - |
| TIL11 | Pollen | 2 Reps | 2 Reps | 2 Reps | - | - | - | - | - | - |
| TIL11 | Root tips | 2 Reps | - | - | - | - | - | - | - | - |
